# A Multiclass Mechanistic Model of Vaccine-Induced Humoral Responses

**DOI:** 10.64898/2026.10.14.759230

**Authors:** Sarafa A. Iyaniwura, Bryan T. Mayer, William O. Hahn, Soo Young Kim, Yunda Huang, Ollivier Hyrien

## Abstract

Vaccine-induced humoral immunity emerges from complex interactions among competing B-cell populations and antibody production in response to antigen exposure, yet the mechanisms governing these interactions remain only partially understood. We developed a mechanistic mathematical model that integrates vaccine regimen features, immunogen kinetics, multiclass B-cell dynamics, and germinal center processes. The model was calibrated to data from the G001 HIV vaccine trial evaluating eOD-GT8, a germline-targeting immunogen designed to activate VRC01-class precursor B cells capable of developing into broadly neutralizing antibody-producing cells. The model reproduced key features of the observed B-cell and antibody responses, enabled estimation of class-specific activation and competition parameters, and supported investigation of the identifiability of competition parameters. Activation rates varied across B-cell classes, with memory B cells exhibiting substantially higher activation rates than naïve B cells. Importantly, competition was stronger within than between classes, with the strongest competition occurring among VRC01-class B cells. These findings suggest that class-specific differences in activation and competition shape the magnitude and composition of vaccine-induced humoral response and may influence the expansion of broadly neutralizing antibody precursors. More broadly, the model provides a quantitative framework for dissecting multiclass B-cell competition and antibody responses, and for evaluating how vaccination strategies shape humoral immunity.

## 1. Introduction

Vaccination is a cornerstone of public health, largely because it confers long-lasting protection through durable humoral responses (1, 2). Following antigen exposure, activated B cells enter germinal centers (GCs), where they undergo clonal expansion, somatic hypermutation, and antigen-driven selection before differentiating into plasma cells and memory B cells (3–5). These interdependent processes span molecular, cellular, and tissue scales, making their coordinated dynamics challenging to characterize *in vivo*, especially since key stages occur in secondary lymphoid tissues that are difficult to observe.

Dynamic models have been used to investigate GC formation, B-cell competition, affinity maturation, the generation of long-lived plasma cell and memory B cells, and antibody production (6–16). These approaches have been applied to vaccination, connecting B-cell dynamics, antigen availability, and antibody kinetics to vaccine-induced humoral immunity. Hodgson et al. (17) linked variability in memory B-cell proliferation to differences in neutralizing antibody responses between SARS-CoV-2 vaccines, while Deng et al. (18) predicted age-dependent heterogeneity in responses to mRNA COVID-19 vaccines. Other studies have examined how antigen kinetics shape B-cell selection and affinity maturation and modeled antibody production and waning following mRNA vaccination (12). More recently, a model integrated B-cell proliferation, plasma-cell differentiation, and memory B-cell generation across vaccine platforms (19).

However, existing models rarely integrate vaccine regimen characteristics with GC dynamics and the downstream generation of memory B cells, plasma cells, and antibodies in a unified mechanistic framework. B-cell development occurs in a competitive environment where different B cells compete for scarce resources. Throughout, we use the term “class” specifically to describe these groups of competing B cells, defined by distinct epitope specificities, affinity classes, or B-cell receptor (BCR) sequence features, as opposed to different B-cell phenotypes (e.g., memory B cells versus plasmablasts) or antibody isotypes (e.g., IgG versus IgM). Despite the central role of such competition in shaping humoral immunity, existing models often do not explicitly represent the dynamics and interactions of multiple B-cell and antibody classes. The inability to characterize responses from different classes in a competitive B-cell environment may be a limitation because, while vaccines are designed to induce protective antibodies against specific regions (“epitopes”) of the vaccine antigen, they contain epitopes that do not contribute to protective immunity but instead elicit competing, “off-target” responses. The competition between B-cell classes can influence the selection, affinity maturation, and immunodominance of desired (“on-target”) B cells, thereby shaping the magnitude, quality, and durability of humoral immunity (16, 20–24). Consequently, failure to account for competition among B-cell classes may limit the ability of current models to characterize vaccine-induced immune responses and quantitatively predict responses to alternative vaccination regimens.

To begin addressing this gap, we develop a deterministic dynamic model of vaccine-induced B-cell and antibody dynamics that links vaccine regimen characteristics, competitive GC processes, and downstream humoral responses. Modern experimental approaches can measure B-cell and antibody responses at phenotypic, epitope-specific, and functional levels. Therefore, the proposed framework represents multiple B-cell classes and integrates their activation, expansion, differentiation, and competition within GCs with immunogen kinetics, the generation of plasma and memory B cells, and antigen-specific antibody production. By combining class-specific competition for immunogen and T follicular helper (T_FH_) cell support with antibody-mediated feedback on immunogen availability, it provides a mechanistic basis for investigating how B-cell interactions shape humoral responses.

Within GCs, these competitive interactions modulate B-cell division and differentiation rates based on immunogen availability and the abundance of competing B-cell classes (22, 25). Competition is allowed to vary across B-cell class pairs, capturing heterogeneity in competitive fitness and resource utilization. Antibody feedback also connects downstream antibody responses to immunogen exposure, allowing assessment of how antibody-mediated changes in immunogen availability influence B-cell selection and expansion following immunization (25).

We apply the model to study B-cell and antibody responses against distinct epitopes, induced by the protein-based eOD-GT8 60mer germline-targeting vaccine evaluated in the International AIDS Vaccine Initiative (IAVI) G001 Phase 1 trial (26). The eOD-GT8 60mer immunogen was specifically designed to engage a clearly defined population of rare VRC01-class precursor B cells, making this study particularly suitable for investigating competition between targeted and non-targeted B-cell populations. By calibrating the model to the G001 data, we estimate class-specific B-cell activation and competition parameters and evaluated how differences in these parameters contribute to the observed responses.

The remainder of this article is organized as follows. Section 2 describes the mechanistic ordinary differential equation (ODE)-based model. Section 3 presents its application to the G001 study and associated analyses. Section 4 discusses the findings, limitations, and potential future applications.

## 2. Methods

### 2.1 Mathematical model

This section presents a general mechanistic model of vaccine-induced B-cell and antibody responses. The model incorporates the transport and immunologic availability of vaccine immunogens across key compartments - the injection site (*I*_*S*_), lymphatic compartment (*I*_*L*_), and GC (*I*_*G*_) (Fig. 1) - which support B-cell activation, selection, proliferation, differentiation, and plasma cell maintenance following vaccination (3, 4). It captures the dynamics of immunogen-specific immune cells, including naïve B cells (*N*), activated naïve B cells (*N*_*a*_), GC B cells (*G*), short-lived antibody secreting cells (ASC) (plasmablasts, *P*_*s*_), long-lived plasma cells (*P*_*l*_), and memory B cells (*M*). It also tracks the production and kinetics of free antibodies (*A*). The model accommodates *K* classes of vaccine-induced B cells and antibodies, which may differ in antigen-binding specificity and affinity. Importantly, it captures competitive interactions across B-cell classes within GCs, reflecting their differential access to immunogen and selection during the GC response.

**Figure 1:**
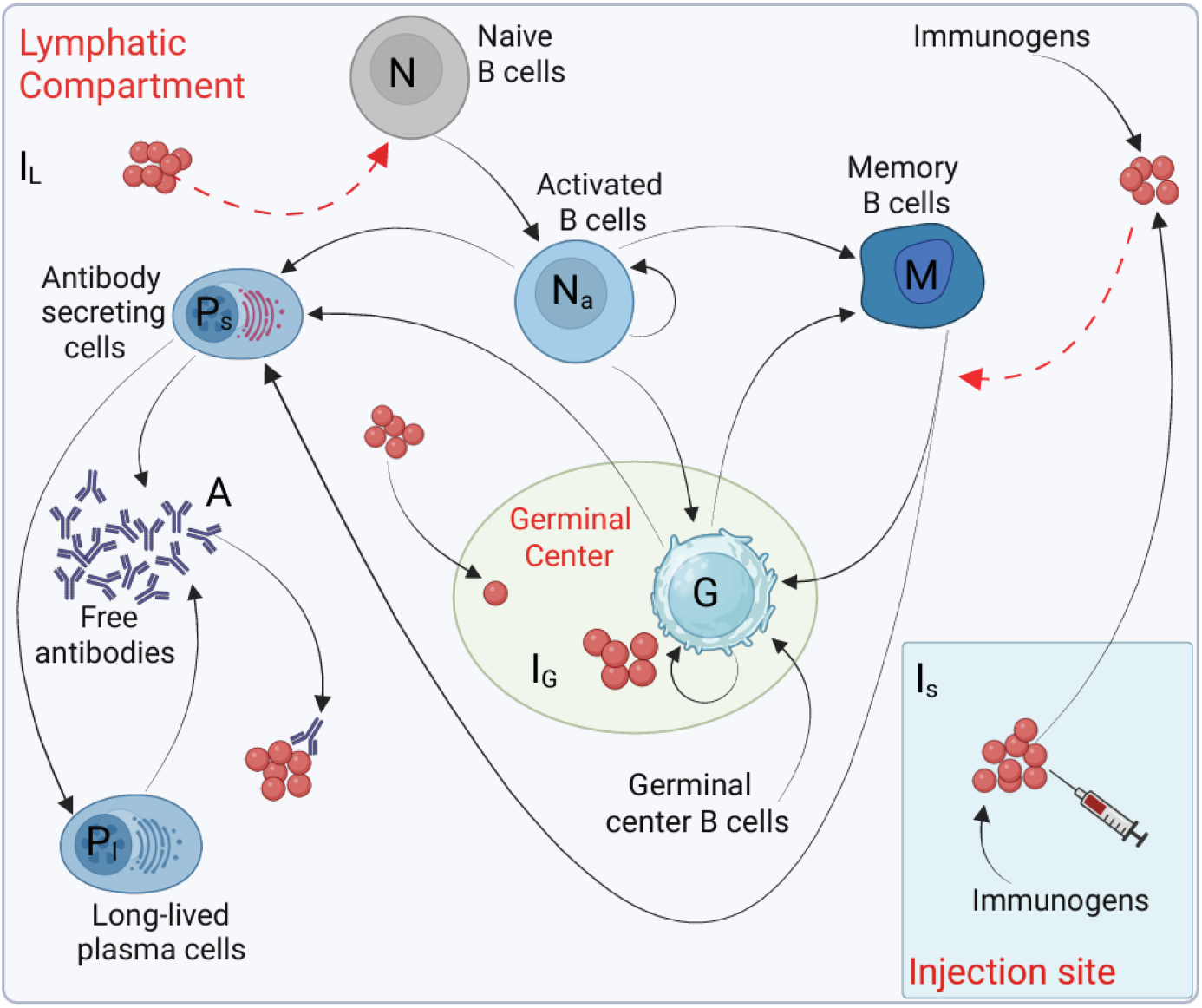
Schematic diagram of B-cell and antibody dynamics in the proposed model. Black arrows indicate immunogen transfer across compartments or B-cell differentiation. Red dashed arrows represent the activation of naïve or memory B cells. *I*_*S*_: immunogen at the injection site, *I*_*L*_: immunogen in the lymphatic compartment, *I*_*G*_: immunogens in the GC, *N*: immunogen-specific naïve B cells, *N*_*a*_: activated naïve B cells, *G*: GC B cells, *P*_*s*_: short-lived ASC, *P*_*l*_: long-lived plasma cells, *M*: memory B cells, and *A*: free antibodies.

**Figure 2.**
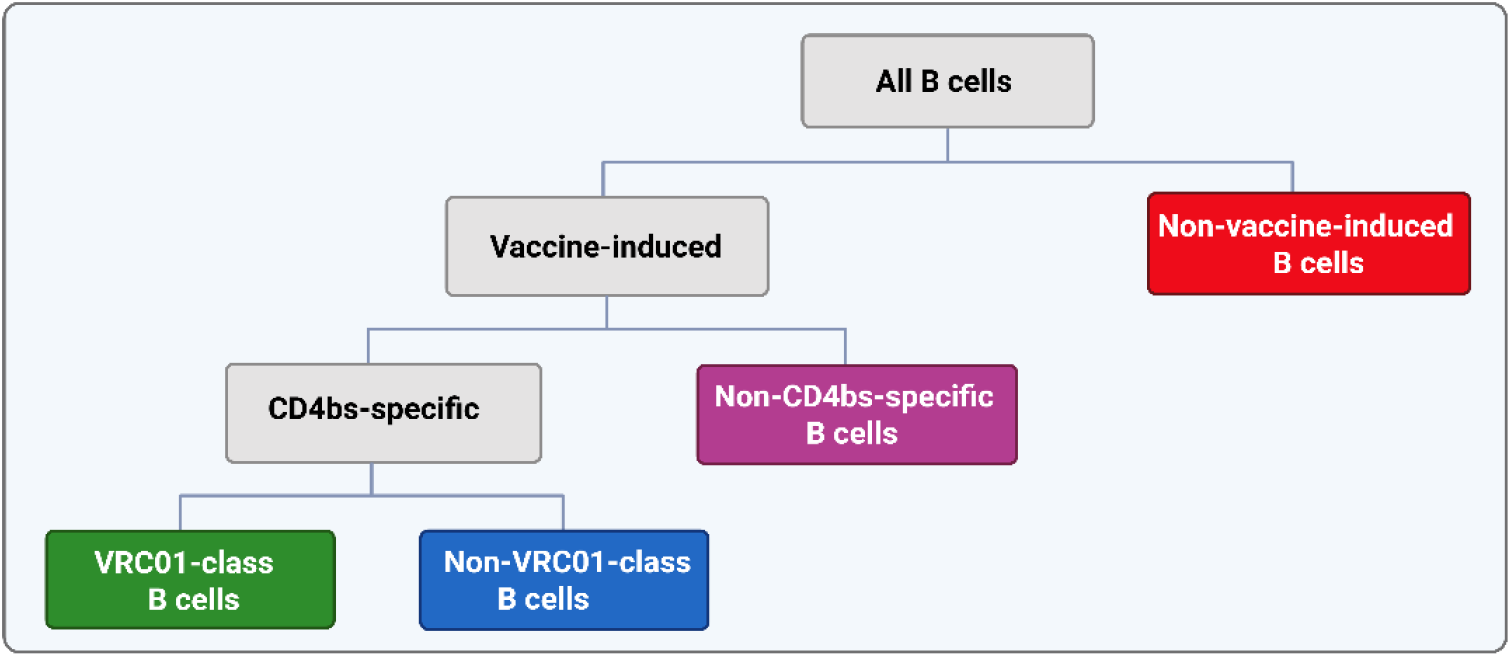
B-cell classes in the G001 model.

Our model shares similarities with those developed by Deng et al. (18), Hodgson et al. (17), and Wilding et al. (19), but extends these frameworks in two key ways. First, it explicitly tracks distinct B-cell classes. Second, it resolves B-cell dynamics within GCs, including immunogen-dependent cell-fate decisions and competition among B-cell classes. In the general model, transitions among B-cell classes can represent changes associated with affinity maturation or altered specificity (27, 28); the G001 model described in Section 3 sets these transition rates to zero, meaning B cells cannot switch between classes.

### 2.2 Vaccine immunogen kinetics

We assume that, upon vaccination, the immunogen is transported from the injection site to the lymph nodes at a rate that depends on factors such as the vaccine platform and delivery route. Mechanistically, examples of these different routes include drainage via afferent lymphatic vessels or transport by antigen-presenting cells (APCs), such as dendritic cells, for protein-based vaccines given either intramuscular or subcutaneous, or production *in situ* by transfected cells followed by transport to lymph nodes for mRNA platforms (29, 30). Within lymph nodes, immunogens are recognized by naïve B cells and presented to T_FH_ cells by antigen-presenting cells, initiating an adaptive immune response (27, 31).

Immunogen kinetics at the injection site are described by

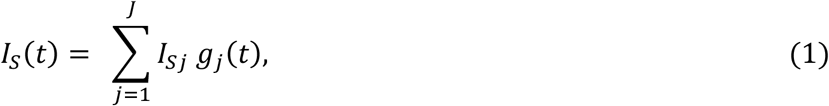

where *I*_*S*_(*t*) denotes the number of immunogen molecules at the injection site, and *t* is the time since administration of the first vaccine dose. Here, *I*_*Sj*_ represents the number of immunogen molecules administered in the *j*^*th*^ vaccine dose at time *t*_*j*_ (*j* = 1, … *J*), and *J* is the total number of doses in the vaccine regimen: *J* = 1 corresponds to a single dose, whereas *J* > 1 represents prime-boost or fractionated dosing regimens (32). The function *g*_*j*_(*t*) describes the fraction of immunogen from dose *j* remaining at the injection site at time *t*, with *g*_*j*_(*t*) = 0, *t* < *t*_*S*_, and *g*_*j*_(*t*) ≥ 0, *t* ≥ *t*_*j*_. In Section 3, we set 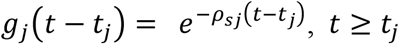, where *ρ*_*sj*_ > 0, *j* = 1, …, *J*, is the immunogen transfer rate from the injection site to the lymphatic compartment (33).

Immunogens in the lymphatic compartment undergo several processes, including degradation at a constant per-capita rate *c*_*l*_, transfer to the GC at rate *ρ*_*l*_ (14), sequestration through binding to circulating antibodies, and engagement of immunogen-specific B cells. The kinetics of immunogens in the lymphatic compartment are therefore described by

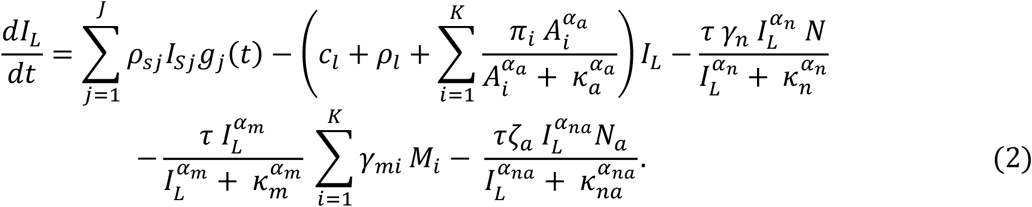

The function 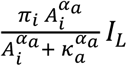 describes the removal of immunogen through binding to vaccine-specific antibodies of class *i* (*A*_*i*_), where *π*_*i*_, *i* = 1, …, *K*, is the corresponding antibody-immunogen binding rate. The functions 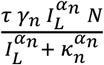 and 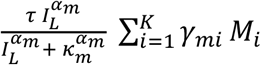 represent immunogen engagement associated with the activation of naïve and memory B cells, respectively. Here, *γ*_*n*_ and *γ*_*mi*_ are the activation rates for naïve B cells (*N*) and memory B cells of class *i* (*M*_*i*_), respectively. The final function represents immunogen engagement associated with proliferation of activated naïve B cells. Here, *τ*(immunogens/cell) is a scaling parameter which characterizes the strength of the immunologic signal required to activate a single naïve or memory B cell. Detailed descriptions of the remaining parameters are provided in Supplementary Material (SM) Table SM1.

Our model describes immunogen kinetics in the GC by

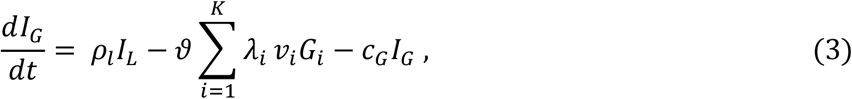

where *I*_*G*_ is the number of immunogen molecules in the GC. Here, *ρ*_*l*_ is the rate of immunogen transfer into the GC, *λ*_*i*_ is the rate at which a GC B cell of class *i* either proliferates, differentiates, or dies, *v*_*i*_ is the probability of division for GC B cell of class *i, c*_*G*_ is the immunogen degradation rate in the GC, and *G*_*i*_ is the size of the class-*i* GC B-cell population.

### 2.3 Naive B-cell dynamics

Following vaccination, vaccine-specific naïve B cells are activated in the lymphatic compartment (34). Their dynamics is governed by

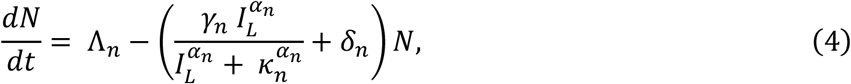

where *N* is the number of vaccine-specific naïve B cells, Λ_*n*_ is the production rate of the naïve B cells, and *δ*_*n*_ is their per-capita death rate. The second term describes activation of naïve B cells.

Following activation, B cells undergo rapid proliferation and clonal expansion, enter GCs, or differentiate into extrafollicular plasmablast or memory B cells (3, 35). The dynamics of activated B cells is described by

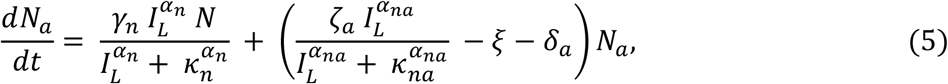

where *N*_*a*_ denotes the number of activated naïve B cells, *ζ*_*a*_ denotes the proliferation rate of activated B cells, and *δ*_*a*_ is their per-capita death rate. The parameter *ξ* represents the rate at which activated B cells commit to one differentiation fate, including GC B cells, extrafollicular short-lived ASCs, or extrafollicular memory B cells. In this equation, the first term represents newly activated naïve B cells, the second term represents proliferation of activated cells, and the final two terms represent the loss of activated cells through differentiation and death, respectively.

### 2.4 Germinal center B-cell dynamics

Recall that *λ*_*i*_ is the rate at which a class-*i* GC B cell undergoes proliferation, differentiation, or death. We first define the probability of division or differentiation, Φ_*i*_(*I*_*G*_, *G*) ∈ [0, 1], as a function of the number of immunogens molecules within the GC, *I*_*G*_, and the number of GC B cells across all classes, *G*= (*G*_1_, *G*_2_, …, *G*_*K*_). The function Φ_*i*_(*I*_*G*_, *G*) is specified to capture immunogen-dependent GC activity and B-cell competition. Here, we set it to

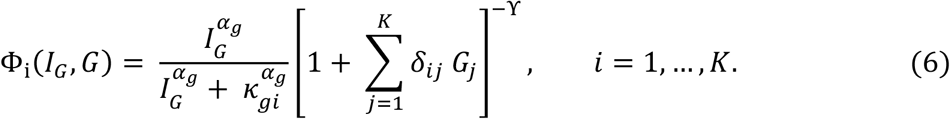

The first term in Eq. (6) describes the dependence of the probability Φ_*i*_(*I*_*G*_, *G*) on immunogen availability. In the absence of immunogen in the GC (*I*_*G*_= 0), this term is null and Φ_*i*_(*I*_*G*_, *G*) = 0, reflecting the biological requirement for immunogen engagement to drive proliferation, somatic hypermutation, and differentiation. As immunogen concentration increases, the term approaches one, indicating saturation, and additional immunogen no longer increases the likelihood of GC progression. The parameter *κ*_*gi*_ is the half-saturation constant governing the influence of immunogen availability on the proliferation and differentiation of class-*i* GC B cells; the Hill coefficient *α*_*g*_ governs the steepness of the relation.

The second term represents competition among GC B cells for immunogen and T_FH_-cell support. B cells that more effectively acquire immunogens and secure T_FH_-cell help gain a competitive advantage over less effective competitors, promoting their survival, proliferation, and maturation (3, 4, 36, 37). The parameter *δ*_*ij*_ represents the strength of competitive suppression experienced by a class-*i* B cell due to a class-*j* B cell. The parameter ϒ ∈ [0,1] modulates the strength of competition, with ϒ = 0 corresponding to a noncompetitive environment and larger values yielding higher competitive pressure. Thus, Eq. (6) couples immunogen availability with population-level competition to describe affinity-dependent selection dynamics within GCs (4).

Given Φ_*i*_ ≡ Φ_*i*_(*I*_*G*_, *G*), the probability that a class-*i* GC B cell divides is

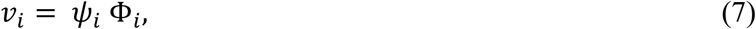

where *ψ*_*i*_ is the conditional probability of division, given that the cell does not die. The probability of differentiation is

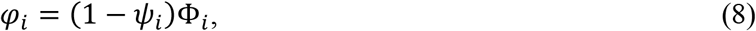

and the probability of cell death is

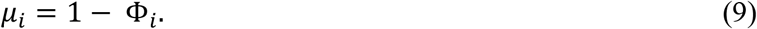

We next describe the population dynamics of GC B cells for the generalized model with *K* B-cell classes. The dynamics account for the entry of naïve and memory B cells into the GC, proliferation within each class, and transitions between classes:

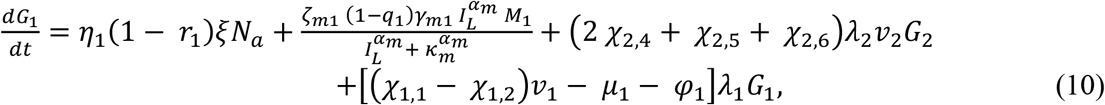

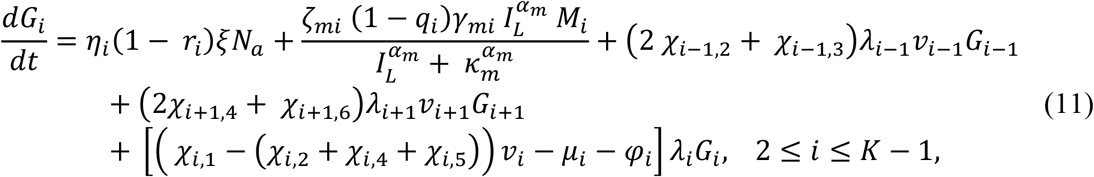

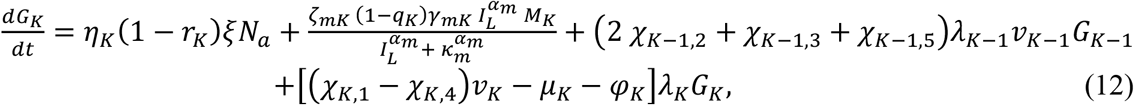

where *η*_*i*_ represents the fraction of activated naïve B cells belonging to class-*i*, and *r*_*i*_ ∈ [0,1] denotes the fraction of activated naïve B cells in class *i* that differentiate into either extrafollicular memory B cells or short-lived ASCs. Thus, 1 − *r*_*i*_ is the fraction that differentiates into GC B cells. Similarly, *q*_*i*_ ∈ [0,1] denotes the fraction of activated class-*i* memory B cells that differentiate into short-lived ASCs, whereas 1 − *q*_*i*_ is the fraction that re-entered the GC. The parameters *χ*_*i,j*_, *j* = 1, … 6, determine how daughter cells distribute across B-cell classes following GC B-cell division (see Table SM1 for details). Finally, *ζ*_*mi*_ is an expansion factor specifying the average number of cells generated by an activated memory B cell before it differentiates or re-enters the GC.

### 2.5 Memory and plasma B-cell dynamics

Following repeated rounds of affinity-based selection within the GC, most B cells fail to survive, while a subset of surviving, affinity-improved B cells exit the GC and differentiate into either antibody-secreting plasmablasts or memory B cells. This process is orchestrated by antigen availability, binding strength, and co-stimulatory signals from T_FH_ cells (3, 4, 37). We incorporate the GC-derived memory B cells and the extrafollicular memory B cells, derived from early activation pathways without entering the GC. Although these two populations may differ due to their developmental origins and potential in their affinity and reactivation (38), their dynamics are described together as a single memory B-cell population within each B-cell class. The temporal evolution of memory B cells is described by

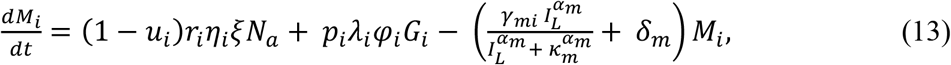

where the first term represents the contribution through the extrafollicular pathway, the second term represents the contribution of GC B cells differentiating into memory B cells upon GC exit, the third term accounts for the loss of memory B cells due to reactivation upon encounter with immunogen, and the last term accounts for death at the per-capita death rate *δ*_*m*_.

In the model, GC-derived plasmablasts, extrafollicular plasmablasts, and plasma cells generated following memory B-cell reactivation are collectively represented as short-lived ASCs. Some of these cells subsequently migrate to the bone marrow where they differentiate into long-lived plasma cells. The dynamics of short-lived ASCs are described by

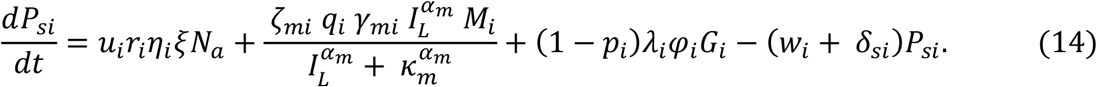

The first term in Eq. (14) accounts for the contribution of extrafollicular plasmablasts to the population of short-lived ASCs, where *u*_*i*_ denotes the fraction of activated class-*i* naïve B cells that bypass the GC and differentiate directly into extrafollicular short-lived ASCs, with the remaining fraction, 1 − *u*_*i*_, differentiating into extrafollicular memory B cells. The second term captures the contribution of memory B cells differentiating into plasma cells, where *q*_*i*_ is the fraction of activated memory B cells differentiating into plasma cells and *ζ*_*mi*_ is the corresponding expansion factor. The term (1 − *p*_*i*_)*λ*_*i*_*φ*_*i*_*G*_*i*_ represents the contribution of GC B cells that exit the GC and differentiate into plasma cells, where *p*_*i*_ represents the fraction of GC B cells of class *i* that differentiate into memory B cells, such that 1 − *p*_*i*_ is the fraction that differentiates into plasma cells. The last term accounts for the loss of short-lived plasma cells through migration to the bone marrow at rate *w*_*i*_ and death at per-capita rate *δ*_*si*_.

Over time, some of the short-lived ASCs migrate to the bone marrow, where they reside in specialized survival niches and differentiate into long-lived plasma cells. These terminally differentiated cells persist for extended periods, ranging from months to years, thereby providing sustained antibody secretion and contributing to long-term humoral immunity (39–41). We describe the dynamics of these long-lived plasma cells by

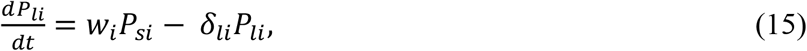

where the first term accounts for the transition of short-lived ASCs to long-lived plasma cells, and the second term represents the loss of these cells due to death at the per-capita death rate *δ*_*li*_.

### 2.6 Antibody dynamics

Lastly, we consider the dynamics of free antibodies, which mediate key effector functions of humoral immunity. Our model explicitly accounts for antibody production by short-lived ASCs and long-lived plasma cells. The kinetics of free antibodies of class *i* (*A*_*i*_) is described by

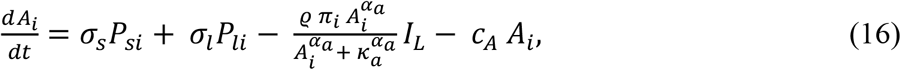

where *σ*_*s*_ and *σ*_*l*_ represent the secretion rates of antibodies by short-lived ASCs and long-lived plasma cells, respectively, while *c*_*A*_ denotes the per-capita elimination rate of circulating antibodies. The function 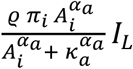 represents antibody removal from the circulation through binding to immunogens in the lymphatic compartment, resulting in sequestration of both free antibodies and immunogens into antibody-immunogen complexes. Here, *π*_*i*_ is the binding rate constant for antibodies of class *i, ϱ* is a scaling parameter accounting for the stoichiometry of antibody-immunogen binding, and *κ*_*a*_ is the half-saturation constant governing the dependence of the binding rate on antibodies concentration. The Hill coefficient *α*_*a*_ captures cooperativity in the binding interaction.

### 2.7 Initial conditions

Together, Eqs. (1)-(16) define the complete model, which describes the dynamics of immunogens, B and plasma cells, and antibodies following immunization for *K* B-cell classes. Detailed descriptions of model variables and parameters are given in Tables 1 and SM1, respectively. The initial conditions at the time of the prime dose are

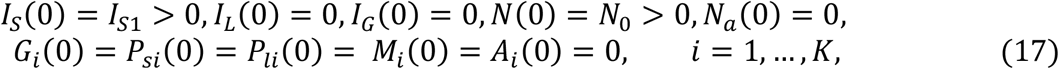

where *I*_*SS*_ > 0, *j* = 1, …, *J*, denotes the number of immunogen molecules in vaccine dose *j*, and *N*_0_ > 0 represents the initial population of vaccine-specific naïve B cells. See Section SM2 for details.

**Table 1.** Model variables and definitions. Some variables (*G, P*_*s*_, *P*_*l*_, *M, A*) are indexed by the B-cell class (*i*) in the model.

| Variable | Description |
| --- | --- |
| $I_S$ | Number of immunogen molecules at the injection site |
| $I_L$ | Number of immunogen molecules in the lymphatic system |
| $I_G$ | Number of immunogen molecules in the GC |
| $N$ | Number of immunogen-specific naïve B cells |
| $N_a$ | Number of activated naïve B cells |
| $G$ | Number of GC B cells |
| $P_s$ | Number of short-lived ASC (plasmablast) |
| $P_l$ | Number of long-lived plasma cells |
| $M$ | Number of memory B cells |
| $A$ | Number of free antibodies |

## 3. Application to the eOD-GT8 germline-targeting vaccine

### 3.1 Biological and clinical context

An effective HIV vaccine has remained elusive partly because the virus can evade immune protection through its extraordinary genetic diversity and rapid evolution (42). A major strategy for overcoming this challenge is to elicit broadly neutralizing antibodies (bnAbs), which can neutralize diverse HIV strains by targeting highly conserved regions of the HIV envelope (Env) that are important for the viral life cycle (43). One prominent Env target is the CD4-binding site (CD4bs), a primary receptor HIV uses to enter the target CD4+ T cell, which is recognized by multiple bnAbs, including VRC01-class bnAbs which are characterized by their use of the VH1-2*02 or VH1-2*04 human heavy chain alleles and a five-amino-acid-long light-chain complementary determining region (CDRL3) (44, 45).

A major challenge for this strategy is that VRC01-class B-cell precursors are rare and generally do not recognize native HIV Env, making their initial activation and maturation challenging. Germline-targeting immunogens address this bottleneck by preferentially activating these precursors. However, following activation, VRC01-class B cells must compete within GCs for resources and survival signals with other VRC01-class B cells, as well as B cells targeting the same CD4bs or other epitopes. This competition may constrain their expansion and maturation, thereby impacting the efficacy of this vaccination strategy.

### 3.2 The IAVI G001 trial

IAVI G001 was a Phase 1, randomized, double-blind, placebo-controlled trial that evaluated the safety and immunogenicity of eOD-GT8 60mer, a recombinant protein HIV vaccine candidate adjuvanted with AS01_B_ (NCT03547245) (26). eOD-GT8 60mer was designed to activate VRC01-class precursor B cells (46, 47). The trial enrolled 48 participants randomized to receive a low-dose vaccine, high-dose vaccine, or placebo. In the low-dose group, 18 participants received 20 *μg* eOD-GT8 60mer with 50 *μg* AS01_B_ and six received placebo; in the high-dose group, 18 participants received 100 *μg* eOD-GT8 60mer with 50 *μg* AS01_B_ and six received placebo. Vaccine or placebo was administered on day 0 and 8 weeks later. Samples were collected at study weeks −4 (baseline), 4, 8, 10, and 16 to measure memory B-cell frequencies and antibody levels (26). The number of days post prime at which measurements were obtained varied across participants.

### 3.3 G001 model specification

To parameterize the model, the two vaccine doses (*J* = 2) were converted from mass to numbers of immunogen molecules, yielding 2.96 × 10^12^ molecules for the low dose (20 *μg*) and 1.48 × 10^13^ molecules for the high dose (100 *μg*) of eOD-GT8 60mer. We restricted the analysis to three relevant B-cell classes: VRC01-class, non-VRC01-class CD4bs-specific, and non-CD4bs-specific B cells. Accordingly, we set *K* = 3. A constant population of non-vaccine-induced B cells was included as background population when calculating memory B-cell frequency for each class (28). Two participants, one in the low-dose group (ID = 28) and one in the high-dose group (ID = 59), had undetectable VRC01-class memory B-cell responses at all timepoints because they lacked the VH1-2*02 and *04 alleles required for VRC01-class responses (48).

The fraction of activated naïve B cells belonging to each of the three B-cell classes (*η*_1_ for non-CD4bs-specific, *η*_2_ for non-VRC01 CD4bs-specific, and *η*_3_ for VRC01-class B cells) were parameterized as

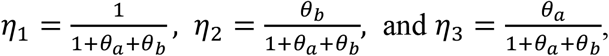

where *θ*_*a*_ > 0 and *θ*_*b*_ > 0 are parameters estimated from the data. We assumed that affinity for the eOD-GT8 60mer immunogen could differ across B-cell classes, with VRC01-class B cells expected to have the highest affinity, followed by non-VRC01 CD4bs-specific and then non-CD4bs-specific B cells (26).

Because these B-cell classes do not communicate, the B-cell dynamics in Eqs. (10–12) simplifies. Specifically, we set *χ*_*i*,1_ = 1 and *χ*_*i,S*_ = 0, *j* = 2, … 6, indicating that B cells of class *i* produce progeny exclusively within the same class. Under these assumptions, Eqs. (10–12) reduce to

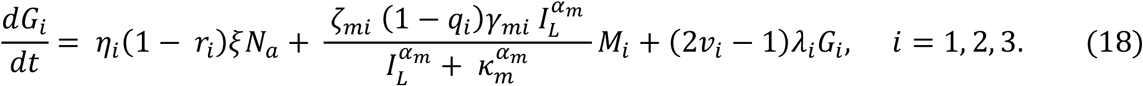

Eq. (18), together with the remaining ODEs (Section SM1), defines the G001 model.

The model predicts the size of plasma cell and memory B-cell populations as well as free antibody levels. Memory B-cell frequencies were computed by dividing the size of each class by that of the total memory B-cell population at each timepoint, with the non-vaccine-induced memory B-cell population held constant at 10^8^ cells, consistent with homeostatic levels in healthy adults (28). Because the model represents B-cell dynamics within a single GC and thus describes the average response of vaccine-induced GCs, model outputs were scaled by an expected number of GCs engaged in the immune response, set to 1,000 (49). The model was fitted to six endpoints (four memory B-cell frequency endpoints and antibody levels against two antigens) measured at five to six timepoints per participant varying across participants.

### 3.4 Model calibration

The model includes a relatively large number of parameters because it represents humoral responses mechanistically across multiple interacting B-cell classes. This level of detail is necessary to distinguish the contributions of different B-cell classes when evaluating how humoral responses may change under alternative vaccination regimens, including different vaccine doses. To calibrate the model, we fixed established biological parameters to values derived from literature and estimated the remaining parameters using data from the G001 clinical trial. Limitations associated with calibration using data from a single clinical trial and opportunities for further model development are discussed in Section 4.2.

#### 3.4.1 Fixed parameter values

The death rate of vaccine-specific naïve B cells was set to 0.011/day (average lifespan ≈ 90 days) (50), with the homeostatic production rate computed as Λ_*n*_ = *δ*_*n*_*N*_0_, where *N*_0_ > 0 is the initial size of the vaccine-specific naïve B-cell population, set to *N*_0_ = 1 × 10^7^ cells (50), ensuring the population remains constant at steady state absent vaccination. The death rate of activated naïve B cells was set to *δ*_*a*_ = 0.33 (average lifespan ≈ 3 days) (50), and that of short-lived antibody-secreting cells (ASCs) at 0.2/day (average lifespan ≈ 5 days) (35, 41, 51). GC B-cell proliferation was set to 4 divisions/day (3, 36, 52). Short- and long-lived plasma cells were assumed to produce antibodies at rates *σ*_*s*_ = *σ*_*l*_ = 2000 antibodies/sec (1.73 × 10^8^ antibodies/day) (44), with free serum antibody degradation set to *c*_*A*_ = 0.041/day (average half-life ≈ 17 days) (53–55).

Hill-type saturation functions describe the effect of immunogen availability on naïve and memory B-cell activation, activated B-cell proliferation, GC B-cell fate, and antibody-immunogen interactions. All Hill coefficients were set to *α*_*a*_ = *α*_*na*_ = *α*_*n*_ = *α*_*m*_ = *α*_*g*_ = 2, yielding a switch-like response to immunogen availability. The half-saturation constants for naïve B-cell activation (*κ*_*n*_), memory B-cell activation (*κ*_*m*_), and activated naïve B-cell proliferation (*κ*_*na*_) were set to *κ*_*n*_ = *κ*_*m*_ = *κ*_*na*_ = 10^6^ immunogens, consistent with Wilding et al.’s estimate (19). The half-saturation constant for the antibody-immunogen interaction was fixed at *κ*_*a*_ = 10^10^. The GC B-cell competition strength was set to ϒ = 0.5. To assess the sensitivity of our results to this assumption, we studied the effect of this parameter on GC B-cell dynamics (Section 3.5.6). The scaling parameters *τ* and *ϑ*, which represent the immunogenic signal required to activate a single naïve or memory B cell, and the signal required for a GC B cell to divide or differentiate, respectively, were both fixed at 1 immunogen/cell. The stoichiometric parameter *ϱ* governing antibody-immunogen binding was fixed at 1, and the antibody-immunogen binding rate was fixed at *π* = 10^−7^/Ab/day.

#### 3.4.2 Estimated parameter values

We estimated the remaining parameters by fitting the G001 model [Eq. (S1) - (S11)] to the G001 data using a nonlinear mixed-effects (NLME) approach and the stochastic approximation expectation–maximization (SAEM) algorithm implemented in Monolix (56, 57). Vaccine dose was evaluated as a covariate on model parameters. Marginal likelihoods were approximated using importance sampling Monte Carlo integration over random effects (35, 36). Model comparisons were performed using log-likelihoods and corrected Bayesian information criteria (BICc) (58). Left-censored measurements were treated as such in the analysis.

The parameters estimated from the G001 data included the activation rates of naïve (*γ*_*n*_) and memory (*γ*_*mi*_) B cells, with *γ*_*mi*_ estimated at the B-cell class level to capture potential differences in memory B-cell reactivation. We also estimated the memory B-cell death rate (*δ*_*mi*_) and the expansion factor *ζ*_*mi*_, representing the average number of cells produced by an activated memory B cell before differentiating into a plasma cell or GC re-entry. For GC dynamics, the half-saturation constant *κ*_*gi*_, *i* = 1, 2, 3, governing the effect of immunogen availability on B-cell division and differentiation was estimated at the B-cell class level, along with the competition coefficients *δ*_*ij*_, *i* = 1, 2, 3 and *j* ≥ *i*, representing the effective competitive suppression of class-*i* B cells by class-*j* B cells in the GC. We assumed *δ*_*ij*_ = 0 for *j* < *i*, to model the assumption that lower-affinity B cells did not competitively suppress higher-affinity B cells within the GC.

### 3.5 Model fit and parameter estimates

#### 3.5.1 Population- and individual-level model-based responses

Parameter estimates are summarized in Table 2. The model reproduced observed memory B-cell frequencies and antibody levels at the population and individual levels in both vaccine groups (Fig. 3). The model reflected biological differences (e.g., genetic capacity for B-cell responses) and vaccination features across participants, predicting absence of VRC01-class responses in two participants (ID=28 and 59) who lacked the VH1-2*02 and VH1-2*04 heavy-chain alleles required for VRC01-class B-cell responses (Figs. SM1, SM3). The model also accounted for the missed boost when estimating the response trajectory of participant ID=172 who missed their second vaccination (Figs. SM3, SM4).

**Table 2.**
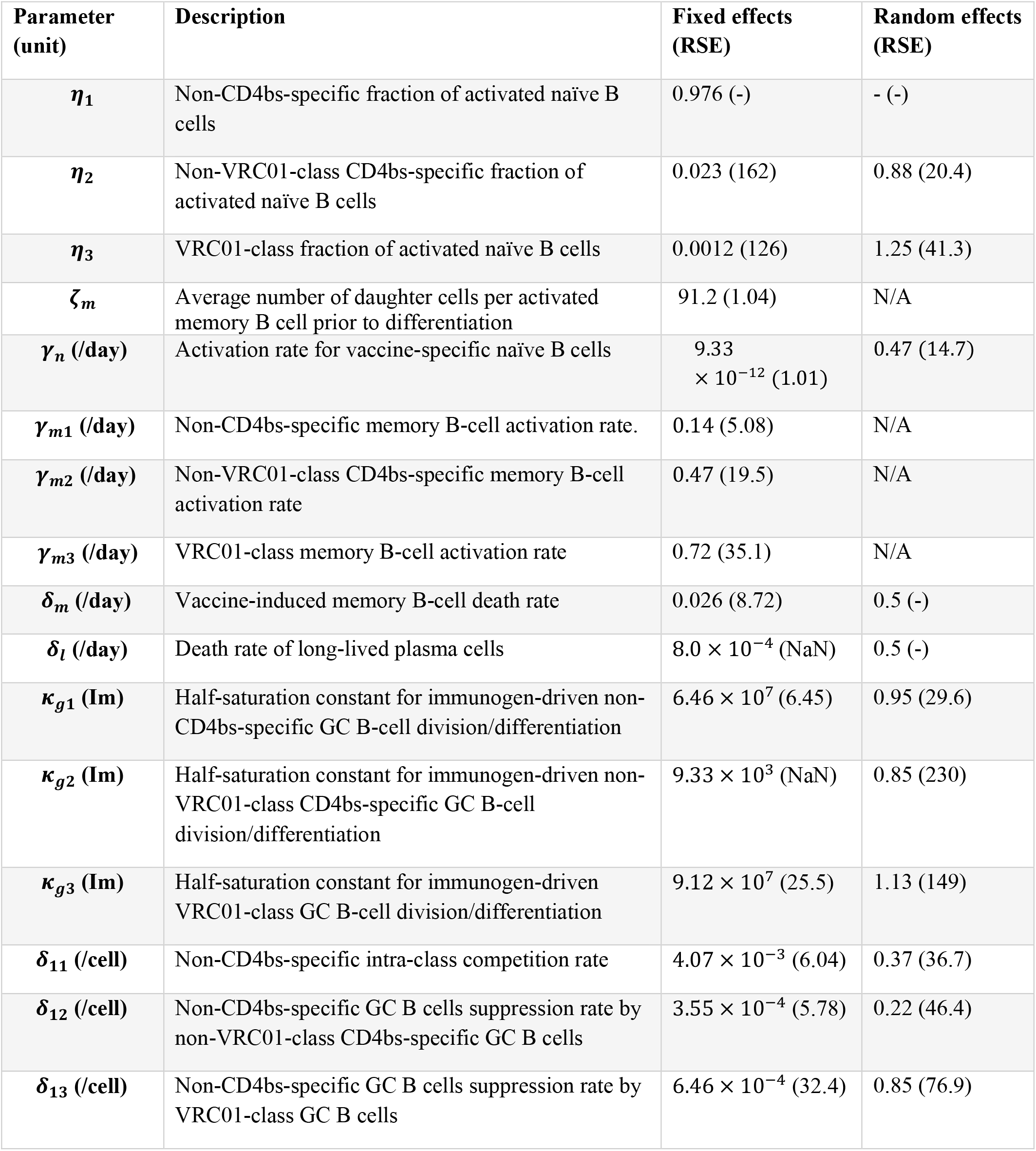
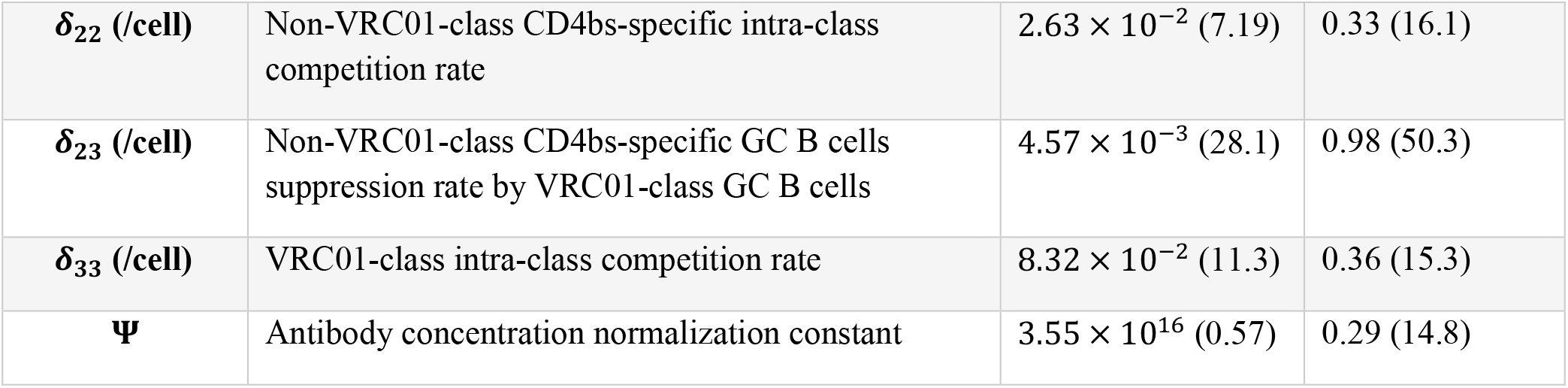
Estimated population-level parameters. RSE is the standard error of the parameter estimate divided by the absolute value of the estimate, expressed as a percentage. A dash (-) indicates that RSE was not applicable for parameters that were fixed or derived from other estimated parameters. “N/A” indicates that no random effect was assumed. “NaN” indicates that the uncertainty in parameter estimate was too large to compute a reliable RSE.

| Parameter (unit) | Description | Fixed effects (RSE) | Random effects (RSE) |
| --- | --- | --- | --- |
| $\eta_1$ | Non-CD4bs-specific fraction of activated naïve B cells | 0.976 (-) | - (-) |
| $\eta_2$ | Non-VRC01-class CD4bs-specific fraction of activated naïve B cells | 0.023 (162) | 0.88 (20.4) |
| $\eta_3$ | VRC01-class fraction of activated naïve B cells | 0.0012 (126) | 1.25 (41.3) |
| $\zeta_m$ | Average number of daughter cells per activated memory B cell prior to differentiation | 91.2 (1.04) | N/A |
| $\gamma_n$ (/day) | Activation rate for vaccine-specific naïve B cells | $9.33 \times 10^{-12}$ (1.01) | 0.47 (14.7) |
| $\gamma_{m1}$ (/day) | Non-CD4bs-specific memory B-cell activation rate. | 0.14 (5.08) | N/A |
| $\gamma_{m2}$ (/day) | Non-VRC01-class CD4bs-specific memory B-cell activation rate | 0.47 (19.5) | N/A |
| $\gamma_{m3}$ (/day) | VRC01-class memory B-cell activation rate | 0.72 (35.1) | N/A |
| $\delta_m$ (/day) | Vaccine-induced memory B-cell death rate | 0.026 (8.72) | 0.5 (-) |
| $\delta_l$ (/day) | Death rate of long-lived plasma cells | $8.0 \times 10^{-4}$ (NaN) | 0.5 (-) |
| $\kappa_{g1}$ (Im) | Half-saturation constant for immunogen-driven non-CD4bs-specific GC B-cell division/differentiation | $6.46 \times 10^7$ (6.45) | 0.95 (29.6) |
| $\kappa_{g2}$ (Im) | Half-saturation constant for immunogen-driven non-VRC01-class CD4bs-specific GC B-cell division/differentiation | $9.33 \times 10^3$ (NaN) | 0.85 (230) |
| $\kappa_{g3}$ (Im) | Half-saturation constant for immunogen-driven VRC01-class GC B-cell division/differentiation | $9.12 \times 10^7$ (25.5) | 1.13 (149) |
| $\delta_{11}$ (/cell) | Non-CD4bs-specific intra-class competition rate | $4.07 \times 10^{-3}$ (6.04) | 0.37 (36.7) |
| $\delta_{12}$ (/cell) | Non-CD4bs-specific GC B cells suppression rate by non-VRC01-class CD4bs-specific GC B cells | $3.55 \times 10^{-4}$ (5.78) | 0.22 (46.4) |
| $\delta_{13}$ (/cell) | Non-CD4bs-specific GC B cells suppression rate by VRC01-class GC B cells | $6.46 \times 10^{-4}$ (32.4) | 0.85 (76.9) |
| $\delta_{22}$ (/cell) | Non-VRC01-class CD4bs-specific intra-class competition rate | $2.63 \times 10^{-2}$ (7.19) | 0.33 (16.1) |
| $\delta_{23}$ (/cell) | Non-VRC01-class CD4bs-specific GC B cells suppression rate by VRC01-class GC B cells | $4.57 \times 10^{-3}$ (28.1) | 0.98 (50.3) |
| $\delta_{33}$ (/cell) | VRC01-class intra-class competition rate | $8.32 \times 10^{-2}$ (11.3) | 0.36 (15.3) |
| $\Psi$ | Antibody concentration normalization constant | $3.55 \times 10^{16}$ (0.57) | 0.29 (14.8) |

**Figure 3.**
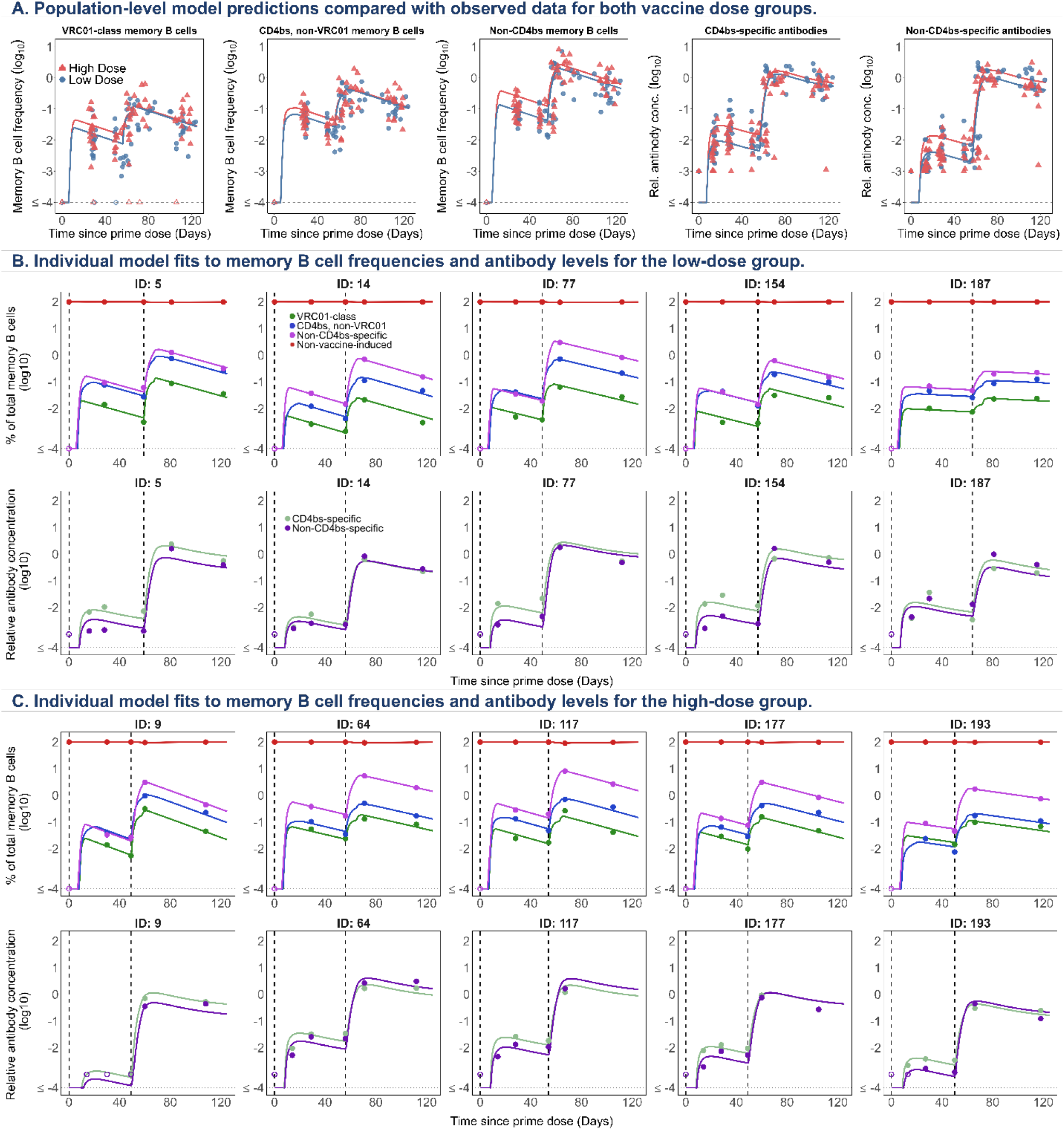
Model fits to memory B-cell and antibody dynamics in the G001 trial. (A) Population-level model predictions for the low-dose (blue) and high-dose (red) groups. Solid lines show model predictions generated using the estimated population-level parameters (Table 2), overlaid on observed data from all G001 trial participants. (B, C) Representative individual-level predicted trajectories for memory B-cell frequencies and antibody levels in the low-dose (B) and high-dose (C) groups. Dots: observed data; solid lines: model predictions; open symbols: observations below the detection threshold. Dashed vertical lines mark the timing of the prime and boost doses. Data are plotted at actual, rather than scheduled, visit days.

#### 3.5.2 Parameters governing immunogen and B-cell dynamics

The rate of immunogen transfer from the injection site to the lymphatic compartment was estimated at *ρ*_*s*1_ = *ρ*_*s*2_ = 2.0/day (half-life ≈ 8 hrs, >99% transfer in 55 hrs), and from the lymphatic compartment to the GC at *ρ*_*l*_ = 2.0/day (half-life ≈ 8 hrs). Immunogen degradation rates in both the lymphatic compartment and GCs were estimated at *c*_*b*_ = *c*_*g*_ = 1.0/day (half-life ≈ 17 hrs), consistent with previous studies of recombinant proteins (34, 59). The proliferation rate of activated naïve B cells was similarly estimated at *ζ*_*a*_ = 2.5/day, while their maturation rate before differentiation into extrafollicular or GC B cells was estimated at *ξ* = 0.45/day (maturation period ≈ 2 days).

The activation rate of vaccine-specific naïve B cells was estimated at 9.33 × 10^−12^/day. Among activated naïve B cells, 0.12% were VRC01-class, 2.25% were non-VRC01-class CD4bs-specific, and 97.6% were non-CD4bs-specific. Activation rates of vaccine-induced memory B cells were estimated separately by B-cell class: VRC01-class memory B cells showed the highest activation rate (0.72/day), followed by non-VRC01-class CD4bs-specific (0.47/day), and non-CD4bs-specific B cells (0.14/day). The higher activation rate of VRC01-class memory B cells is consistent with their preferential responsiveness to the immunogen.

Outside GCs, any activated memory B cell produced, on average, *ζ*_*m*_ ~ 91 progeny before differentiating into a short-lived ASC or re-entering a GC, equivalent to at least 7 cell cycles. The death rate of vaccine-induced memory B cells was estimated at *δ*_*m*_ = 0.026/day, corresponding to an average lifespan of ~ 38.4 days, while the death rate of long-lived plasma cells was estimated at *δ*_*l*_ = 0.0008/day, corresponding to a lifespan of ~3.42 years.

#### 3.5.3 Immunogen sensitivity

Class-specific half-saturation constants controlling the effect of immunogen availability on GC B-cell division and differentiation [Eq. (6)] were lowest for non-VRC01 CD4bs-specific B cells (*κ*_*g*2_ = 9.33 × 10^3^) roughly four orders of magnitude below the estimates for non-CD4bs-specific (*κ*_*g*1_ = 6.46 × 10^7^) and VRC01-class (*κ*_*g*3_ = 9.12 × 10^7^) B cells. A higher half-saturation constant means that more immunogen is needed to drive division and differentiation. Therefore, VRC01-class B cells were the least sensitive to immunogen availability, while non-VRC01 CD4bs-specific B cells responded even with low levels of antigen. Consequently, as immunogen is cleared from the GC, VRC01-class B cells would be the first to fall below the level needed to sustain division and differentiation, a disadvantage relative to the other two classes (Fig. SM6-7). This suggests that maintaining immunogen availability in the GC may be important for keeping VRC01-class B cells competitive long enough to undergo the affinity maturation required for broadly neutralizing activity (60–62).

#### 3.5.4 Effect of vaccine dose

The high-dose group showed modestly higher humoral responses than the low-dose group (Fig. 3A). To determine whether these differences were attributable to dose-dependent changes in model parameters, we evaluated vaccine dose as a covariate on the estimated parameters. We found no evidence for dose-dependent effects on the estimated parameters. Thus, within the structure of the model, the modestly higher responses in the high-dose group were explained by the greater immunogen input and resulting differences in immunogen availability, rather than by dose-dependent changes in the underlying B-cell or antibody dynamics.

#### 3.5.5 Germinal center dynamics

The calibrated model was used to predict GC dynamics for the three B-cell classes in both vaccine groups (Fig. 4). Following both vaccinations, activated naïve B cells entered GCs, with entry peaking within the first two weeks (Fig. 4A). Non-CD4bs-specific B cells contributed the largest number of GC-entering activated B cells, followed by non-VRC01-class CD4bs-specific and VRC01-class B cells, consistent with the estimated fractions of activated naïve B cells reported in Section 3.3. More activated naïve B cells entered the GC in the high-dose than in the low-dose group, reflecting the greater immunogen input. Activated memory B cells similarly entered the GC after each vaccination (Fig. 4B), with entry ~203-fold higher after the boost than after the prime in the low-dose group, and ~87-fold higher in the high-dose group.

**Figure 4.**
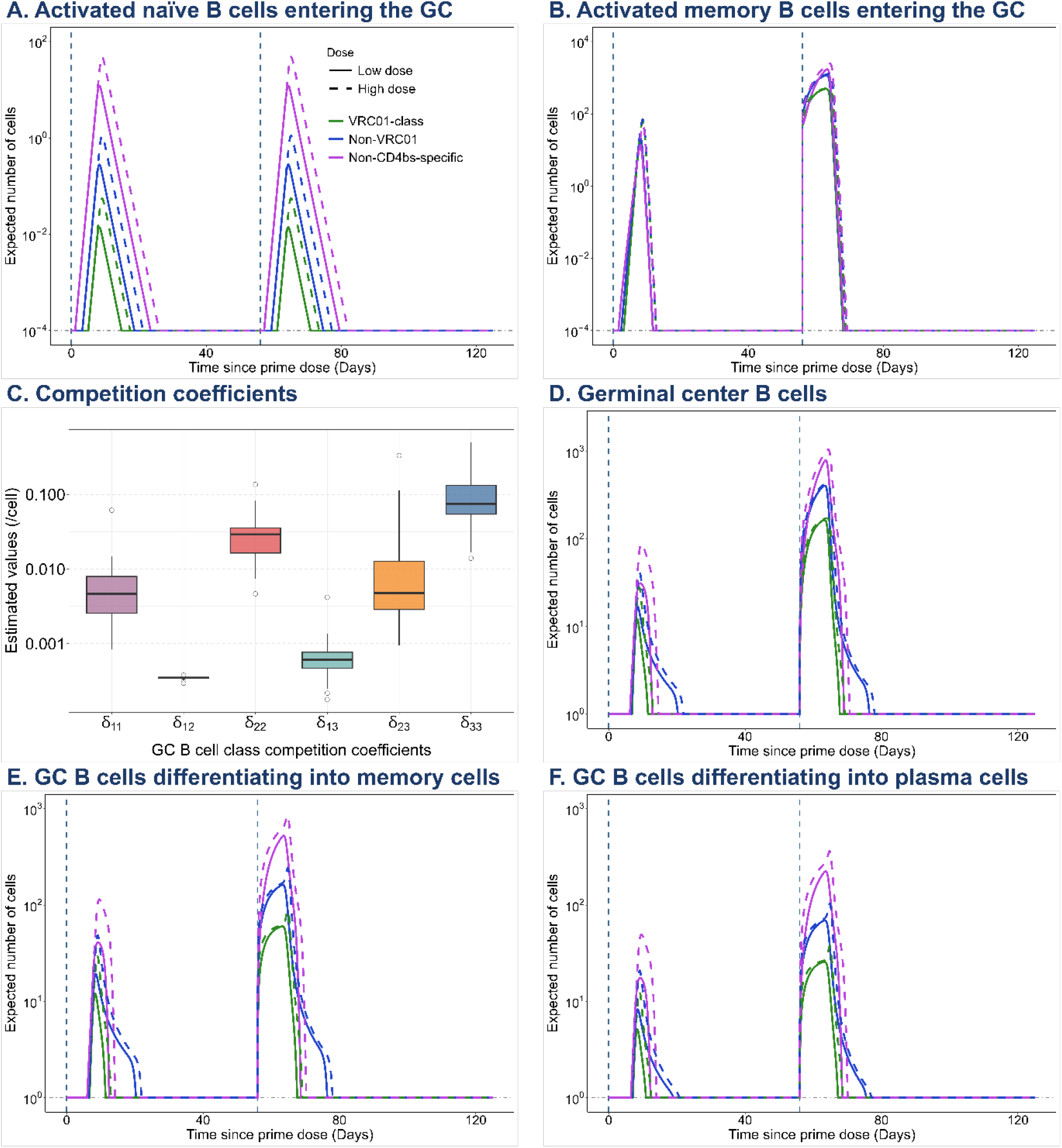
Predictions of GC B-cell dynamics for the low- (solid lines) and high-dose (dashed lines) groups. (A) Activated naïve B cells entering the GC for VRC01-class (green), non-VRC01 CD4bs-specific (blue), and non-CD4bs-specific (purple) B cells. (B) Activated memory B cells entering the GC, by B-cell classes. (C) Distribution of estimated individual GC B-cell competition coefficients (*δ*_*ij*_), by classes. (D) GC B-cell population dynamics, by class. (E) Number of GC B cells differentiating into memory B cells, by class. (F) Number of GC B cells differentiating into plasma cells, by class. In all panels, vertical dashed lines indicate prime (Day 0) and boost (Day 56) vaccination.

The total GC B-cell population (Fig. 4D) expanded sharply after each vaccine dose, with a brief delay after the prime and a larger response following the boost. The higher dose consistently produced a larger and more sustained GC response than the lower dose, and the non-CD4bs-specific class dominated the GC B-cell population throughout the immune response, followed by non-VRC01-class CD4bs-specific and VRC01-class B cells. The dynamics of GC B cells differentiating into memory B cells and plasma cells upon GC exit (Figs. 4E and 4F, respectively) closely mirrored those of the total GC B-cell population, with peak differentiation coinciding with the peak GC response after each dose. The boost dose produced a substantially larger differentiation pulse than the prime dose, consistent with the enhanced GC response driven by memory B-cell reactivation. Across both differentiation pathways, the non-CD4bs-specific B-cell class consistently produced the largest output while VRC01-class B cells contributed the smallest.

#### 3.5.6 Class-specific competition within germinal centers

Within GCs, VRC01-class, non-VRC01-class CD4bs-specific, and non-CD4bs-specific B cells compete for limited access to follicular dendritic cells and T_FH_ cells, which together regulate the probability that a GC B cell divides, differentiates, or dies. This competition was represented by class-specific competition coefficients [Eq. (6)]. The estimated GC B-cell competition coefficients (Fig. 4C) suggest heterogeneity in both within- and between-class competition. The within-class competition coefficient *δ*_33_ (~ 8.32 × 10^−2^/cell) for VRC01-class B cells was the largest estimated coefficient, reflecting that competition was strongest among VRC01-class B cells themselves. The within-class coefficient for non-VRC01-class CD4bs-specific B cells, *δ*_22_ (~ 2.63 × 10^−2^/cell), was also relatively large, whereas *δ*_11_ (~ 4.07 × 10^−3^/cell) for non-CD4bs-specific B cells was the lowest of the three within-class coefficients.

The estimated between-class coefficients were smaller than the corresponding within-class coefficients. In particular, *δ*_13_ (~ 6.46 × 10^−4^/cell), which represents the effect of VRC01-class B cells on non-CD4bs-specific B cells, was lower than *δ*_23_ (~ 4.57 × 10^−3^/cell), representing the effect of VRC01-class B cells on non-VRC01-class CD4bs-specific B cells; both were lower than *δ*_33_, suggesting that VRC01-class B cells were more competitive among themselves than with other B-cell classes. Similarly, *δ*_12_ (~ 3.55 × 10^−4^/cell), representing the effect of non-VRC01-class CD4bs-specific B cells on non-CD4bs-specific B cells, was smaller than *δ*_22_, indicating stronger within-than between-class competition for this B-cell class as well. Overall, these findings indicate that competitive effects were strongest within B-cell classes, particularly among VRC01-class B cells, while weaker but measurable competitive interactions persisted between distinct B-cell classes.

Because the competition parameters play a central role in the model, we next investigated whether they can be uniquely inferred from model trajectories and from the G001 data.

Suppose first that all model parameters are fixed, except for *δ*_*ij*_, *i, j* = 1, …,3. Let *δ* = *δ*_*ij*_, *i, j* = 1, …,3 and 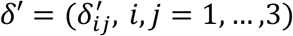. Theorem 1 establishes that, when min*α*_*m*_, 2*α*_*g*_ > 1 and *λ*_1_, *λ*_2_, *λ*_3_ are pairwise distinct, the models parameterized by *δ* and *δ*′ produce the same solution to the G001 Model [(S1)–(S11)] if and only if *δ* = *δ*′. Thus, the competition parameters *δ*_*ij*_ are structurally identifiable under these sufficient conditions. The proof is given in Section SM5. Structural identifiability of the complete parameter set is beyond the scope of this paper.

##### Theorem 1.

*Suppose that all model parameters other than δ are fixed, and that ψ*_*i*_ > 0, *i* = 1, 2, 3, min*α*_*m*_, 2*α*_*g*_ > 1, *and λ*_1_, *λ*_2_, *λ*_3_ *are pairwise distinct. Then, the G001 models parametrized by δ and δ*′ *produce the same solution to the ODEs (S1)–(S11) on any* [0, *T*], *with T* > 0, *if and only if δ* = *δ*′.

**Proof:** Section SM5.

The assumptions in Theorem 1 are sufficient, but not necessary, for identifiability. In particular, to reduce the number of parameters estimated from the G001 data, we assumed *λ*_1_ = *λ*_2_ = *λ*_3_ during model fitting. Consequently, Theorem 1 does not directly establish identifiability for the fitted model but demonstrates that the competition coefficients are structurally identifiable in a closely related setting. We therefore further assessed identifiability and estimability of the competition coefficients under the fitted model using a profile-likelihood analysis (Fig. SM5). The resulting profiles support practical identifiability of the competition coefficients from the G001 data alone. The competition parameters were estimated with good precision, although uncertainty was greater for some between-class than within-class coefficients (Table 2). Within-class coefficients *δ*_11_, *δ*_22_, and *δ*_33_ had RSEs of 6.0%, 7.2%, and 11.3%, respectively, whereas the RSEs for the between-class coefficients ranged from 5.8% to 32.4% (Table 2).

Biologically, the stronger within-class competition observed for VRC01-class B cells may reflect greater overlap in the antigenic and cellular resources required for GC selection. Because VRC01-class B cells recognize the same or overlapping epitopes associated with the HIV Env CD4-binding site, they may compete more directly for access to these epitopes and for T_FH_-cell help. Strong within-class competition may act as a regulatory mechanism that limits the expansion of individual B-cell classes and associated antibodies. By contrast, B cells from different classes may recognize distinct or less-overlapping epitopes and therefore experience less direct competition for antigen. Nevertheless, between-class competition is expected because all B-cell classes share the same GC environment and compete for common limiting resources, particularly T_FH_-cell help, as well as access to antigen and other survival and proliferative signals. Thus, the weaker between-class coefficients may reflect reduced overlap in antigenic resources rather than an absence of competition between B-cell classes.

To characterize how the competition-strength parameter (ϒ) influences GC B-cell dynamics, we simulated the model across a range of ϒ values using the estimated population parameters in Table 2 (Figs. SM6 and SM7). Across all three B-cell classes and in both dose groups, the probability of division and differentiation, *Φ*_*i*_, decreased as the GC B-cell population expanded following vaccination, with the decline becoming more pronounced at larger values of ϒ. These results indicate stronger competitive suppression of GC B-cell expansion as GC activity increases until the GC B-cell population becomes minimal, and converging to zero as GC immunogen level decreases. Although the qualitative behavior was similar across dose groups, the magnitude of the response differed, suggesting that vaccine dose modulates the sensitivity of GC dynamics to competition strength. To evaluate the effect of ϒ on the estimated competition coefficients, we refit the model to the G001 data across the same range of ϒ values and re-estimated the competition parameters (Fig. SM8). Across this range, within-class competition coefficients remained larger than their corresponding between-class coefficients, indicating that the primary conclusion of stronger within-class competition (Table 2) is robust to the assumed value of ϒ.

#### 3.5.7 Antibody-mediated feedback and sensitivity to the antibody-immunogen binding rate

The model represents antibody-mediated feedback by allowing antibodies to bind and remove immunogen from the lymphatic compartment, thereby reducing immunogen availability for B-cell activation and downstream GC responses (25, 63). Because the antibody-immunogen binding-rate parameter *π* could not be reliably estimated from the G001 data, we evaluated predictions across fixed values from *π* = 10^−7^ to 10, together with a no-masking scenario (*π* = 0). In addition, we computed the ratio, relative to baseline (*π* = 10^−7^), of model-predicted immunogen levels (lymphatic compartment and GC), GC B cells, memory B cells, plasma cells, and antibodies across the different values of π, shown in Section SM7 for the high-dose group. The results are similar for the low-dose group.

Masking occurred shortly after vaccination and was most consequential after the boost (~day 56), when antibodies elicited by the prime were present. Increasing *π* produced progressively greater depletion of lymphatic and GC immunogen. For *π* ≤ 10^−3^, immunogen, GC B-cell, memory-B-cell, plasma-cell, and antibody trajectories remained close to the no-masking scenario. At *π* = 10^−2^ and 10^−1^, reductions were modest; at *π* = 10, masking substantially depleted free immunogen after both doses and reduced GC B cells, memory B cells, plasma cells, and free antibodies across all three B-cell classes.

The limited identifiability of *π* is consistent with its predicted effect being restricted to a short post-dose interval that was not sampled in G001. Across refits with fixed *π*, most parameter estimates and the primary conclusions, including heterogeneous B-cell activation and stronger within-class than between-class GC competition, remain stable. The initial activated-naïve-B-cell fractions (*θ*_*a*_ and *θ*_*b*_) showed the greatest relative variation, and selected GC immunogen-sensitivity and competition parameters were uncertain in individual fits (Table SM2).

## 4. Discussion

### 4.1 Mechanistic insights into humoral responses

We developed a mechanistic model of vaccine-induced humoral responses that links vaccine regimen characteristics and immunogen kinetics to competition among B-cell classes, GC processes, and antibody production. The model accommodates multiple B-cell classes, explicitly represents immunogen availability, and captures GC competition. We applied it to characterize humoral responses elicited by the HIV eOD-GT8 60mer immunogen administered as a protein vaccine in antigen-naïve adults.

A key model feature is its explicit representation of competition between each pair of B-cell classes in the GC. This feature, which enables quantitative estimation and analysis of how competition among B-cell classes shapes B-cell selection and expansion, is particularly relevant for vaccines that aim to preferentially activate, expand, and mature specific B-cell populations, such as germline-targeting and sequential immunization strategies.

Our G001 analysis suggested that, although VRC01-class precursor B cells were rare within the activated naïve repertoire, they exhibited strong responsiveness following vaccination, consistent with their preferential engagement by eOD-GT8. These findings suggest that a low precursor frequency does not preclude robust expansion if cells have a competitive advantage relative to other B-cell classes. The model also identified heterogeneous competitive interactions among B-cell classes, with stronger within-than between-class competition, particularly among VRC01-class B cells. This pattern may reflect greater overlap in the antigenic and cellular resources required by B cells within the same class, whereas between-class competition may arise from shared dependence on limiting resources such as T_FH_-cell help. Such a mechanism is consistent with how T-cell help is known to occur (e.g., via presentation of epitopes not recognized by the antibody encoded by a given B cell).

Additionally, memory B-cell activation rates were substantially higher than those of naïve B cells across B-cell classes, consistent with the established property of immunological memory whereby prior antigen exposure enables a more rapid and robust response upon re-exposure.

#### 4.2 Limitations and future directions

Some limitations should be noted. Although antibody-mediated feedback on immunogen availability is represented in the model, the G001 data did not permit precise estimation of the antibody-immunogen binding rate or a definitive assessment of the magnitude of antibody-mediated masking. Additional early post-vaccination sampling, direct measurements of immunogen concentrations or antibody-immunogen complexes, and data from alternative vaccination regimens could improve identifiability of antibody-feedback parameters and clarify the role of antibody-mediated regulation in shaping B-cell dynamics. The current model does not explicitly represent APC, T_FH_-cell, or GC spatial dynamics across the light and dark zones, and model calibration was based only on data from the G001 trial. In addition, B cells in our G001 model do not transition between specificity classes: each cell remained in its assigned class, although class sizes changed over time through activation, expansion, differentiation, and death. This assumption is likely appropriate for describing early vaccine-induced responses, but it does not account for the gradual evolution of affinity and specificity that can occur during affinity maturation.

Future extensions will incorporate explicit APC and T_FH_-cell dynamics, affinity-class transitions, and data from multiple past and ongoing clinical trials evaluating similar HIV vaccine strategies. These extensions will enable more rigorous model calibration and validation, facilitate assessment of model generalizability across studies, and support refinement of the model structure. The framework may further enable *in silico* optimization of vaccine dosing and scheduling. More broadly, such models provide a quantitative foundation for studying how immunization strategies shape B-cell selection and antibody responses, thereby supporting rational vaccine regimen design.

## Data availability statement

The clinical data from the IAVI G001 trial analyzed in this study were previously reported in Leggat et al., (26). The model code and parameter estimates generated during the current study are available from the corresponding author on reasonable request.

## Acknowledgements

We thank the G001 study participants for their time, commitment, and contribution to this research. We are also grateful to the G001 study investigators, clinical staff, laboratory personnel, and study coordinators whose efforts made the collection and generation of these data possible.

## Supplementary Materials

### SM1. The G001 model

Immunogen dynamics in the injection site:

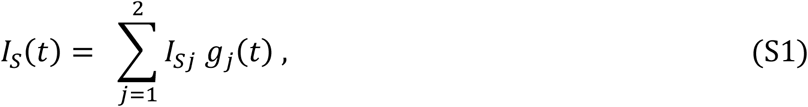

where *g*_*S*_(*t*) = 0, *t* < *t*_*S*_ and 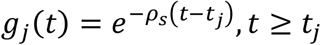, with *ρ*_*s*_ > 0, *j* = 1, 2. Immunogen dynamics in the lymphatic compartment:

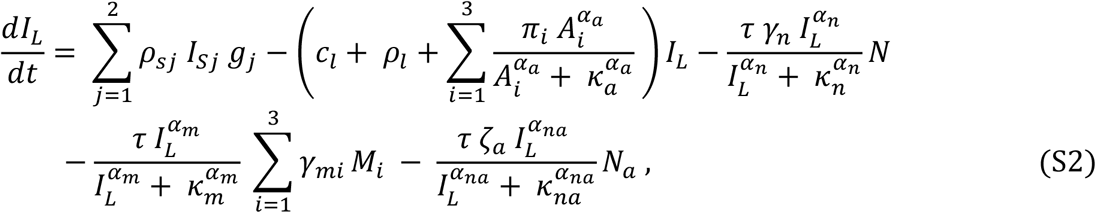

Immunogen dynamics in the germinal center:

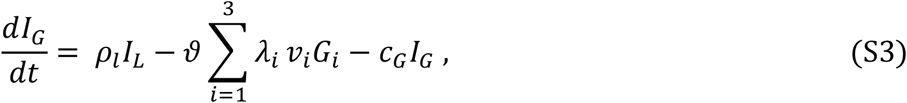

Vaccine-specific naïve B-cell dynamics:

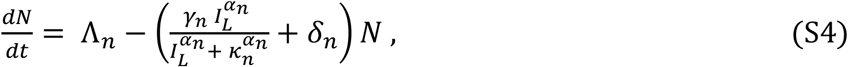

Activated-naïve B-cell dynamics:

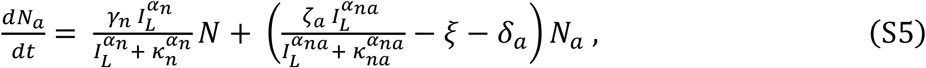

Germinal center B-cell dynamics:

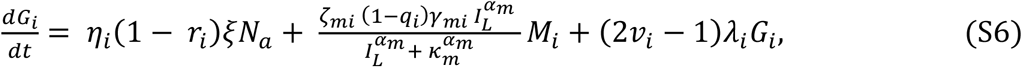

where *v*_*i*_ = *ψ*_*i*_*Φ*_*i*_(*I*_*G*_, *G*), *ψ*_*i*_ > 0, and

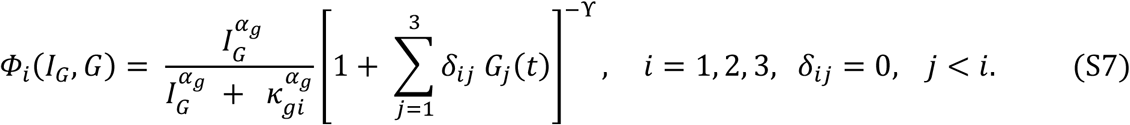

Short-lived antibody secreting cells dynamics:

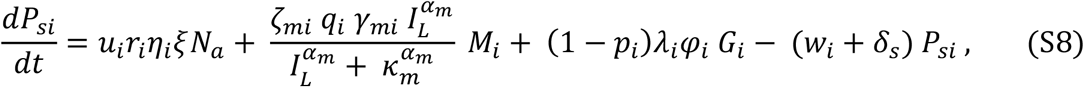

Long-lived plasma B-cell dynamics:

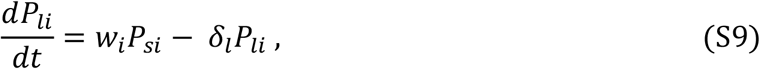

Memory B-cell dynamics:

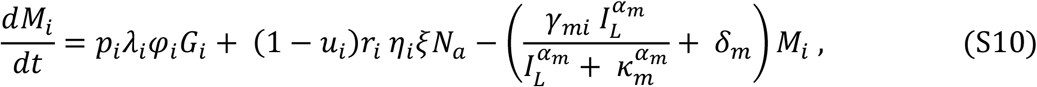

Free antibody dynamics:

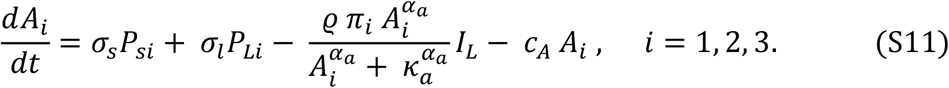

### SM2. Model initial conditions

The ODE model in Eq. (S1)–(S11) describes the dynamics of B cells and antibodies induced by a protein-based immunogen and can be used to simulate both bolus and fractionated prime-boost vaccination regimens by specifying appropriate initial conditions. Here, we describe the initial conditions for both vaccination scenarios in the context of a clinical trial.

Vaccine doses in clinical trials are typically specified in terms of mass (grams); however, our model requires the number of immunogen molecules. Conversion from mass to molecule count is therefore necessary and is detailed in Section SM3. For the G001 trial consisting of two vaccine doses, at the injection site, the prime dose is specified as the initial condition *I*_*S*_(0) = *I*_*S*1_ > 0. The subsequent dose is given by *I*_*S*_(*t*_2_) = *I*_*S*2_ > 0, where *t*_2_ > 0 is the administration time for the second dose. Since this is a homologous vaccine regimen, and the same amount (in grams) of vaccine was administered for both doses, we set *I*_*S*1_ = *I*_*S*2_.

Since the model is applied to antigen-naïve individuals, i.e., those with no prior exposure to the immunogen, all immunogen and B-cell compartments are empty at the time of prime dose administration, except the injection site and vaccine-specific naïve B cells. Accordingly, the initial conditions for immunogens in the lymphatic compartments and GC are *I*_*L*_(0) = 0 and *I*_*G*_(0) = 0, respectively. For the naïve B cell population, we assume a positive constant initial population of vaccine-specific naïve B cells, *N*(0) = *N*_0_ > 0. Since no vaccine-induced immune response exists prior to the prime dose, the initial populations of activated B cells, GC B cells, short-lived ASCs, long-lived plasma cells, and memory B cells are all set to zero, i.e., *N*_*a*_(0) = *G*_*i*_(0) = *P*_*si*_(0) = *P*_*li*_(0) = *M*_*i*_(0) = 0, respectively. Similarly, there are no vaccine-induced antibodies at baseline, so *A*_*i*_(0) = 0 for *i* = 1, 2, 3.

Putting these conditions together, we present the complete initial conditions for the model at the time of the prime dose:

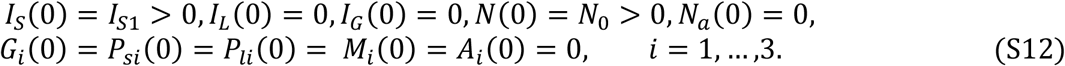

### SM3. Conversion of vaccine dose from mass to molecule count and antibody MFI to relative concentration

In the G001 clinical trial, vaccine doses were specified in terms of physical mass (micrograms, *μg*). However, our model tracks the number of immunogen molecules rather than mass. To convert the vaccine dose (in *μg*) to the number of immunogen molecules, we use the formula

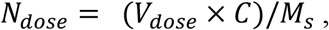

where *N*_*dose*_ is the number of immunogen molecules in a vaccine dose *V*_*dose*_, and *M*_*s*_ is the molecular mass of a single immunogen molecule (in Daltons), and *C* = 6.024 × 10^17^ Daltons/*μg* is the conversion factor from micrograms to Daltons, obtained from the relation 1 *μg* = 6.024 × 10^17^ Daltons. Since the eOD-GT8 60mer immunogen is a protein-based nanoparticle consisting of 60 identical subunits, each with a monomer mass of 67.7 × 10^3^ Daltons, we set *M*_*s*_ = 60 × 67.7 × 10^3^ = 4.062 × 10^6^ Daltons as the molecular mass of a single immunogen nanoparticle. Using this formula, we obtain *N*_*dose*_ = 2.96 × 10^12^ molecules for the 20 *μg* dose and *N*_*dose*_ = 1.48 × 10^13^ molecules for the 100 *μg* dose.

Furthermore, we estimated the relative antibody concentrations for each participant from the MFI measurement in the clinical trial data using a novel statistical method that integrates net MFI values across all available dilutions via quadratic B-splines with monotone constraints. The function describes the relationship between MFI and the log-transformed concentration of binding antibodies in each sample, with the monotone constraints ensuring a biologically consistent, non-decreasing relationship between the two quantities. The model was fitted to the data using the Newton-Raphson procedure, yielding estimates of relative antibody concentrations, i.e., unknown concentrations proportional to a common multiplicative factor shared across all samples. Antibody concentration ratios were subsequently derived from these relative estimates. For example, the percentage of epitope-specific (*P*_*S*_) antibodies were computed as:

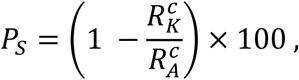

where 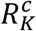 and 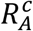 refer to the relative concentration of vaccine-specific antibodies and the relative concentration of vaccine-specific antibodies binding to the mutated CD4bs KO immunogen, respectively. The calculations assumed that *C*_*A*_ = *C*_*K*_ + *C*_*S*_, where *C*_*A*_, *C*_*K*_, and *C*_*S*_ refer to the concentration of antigen-specific, CD4bs-KO, and CD4bs-specific antibodies that bind to the CD4bs.

### SM4. Model fit to the G001 data

**Figure SM1:**
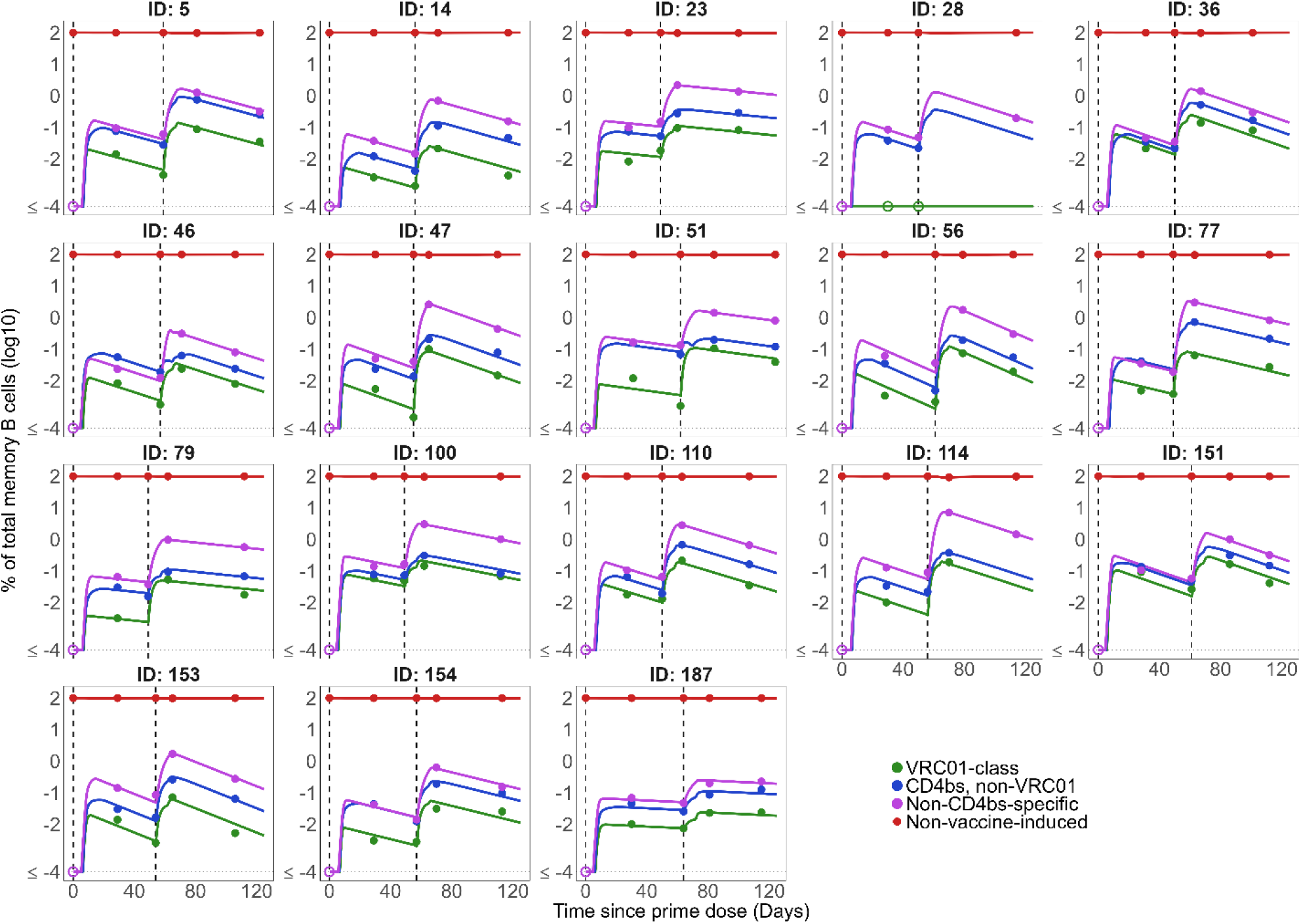
Individual-level predictions for memory B-cell frequencies in the low-dose group. Observed (dots) and model-predicted (solid lines) frequencies of four memory B cell classes (log_10_) as a fraction of total memory B cells, for participants in the low-dose group of the G001 trial. The four classes are: VRC01-class (green), non-VRC01-class CD4bs-specific (blue), non-CD4bs-specific (purple), and non-vaccine-induced (red). Vertical dashed lines indicate vaccinations.

**Figure SM2:**
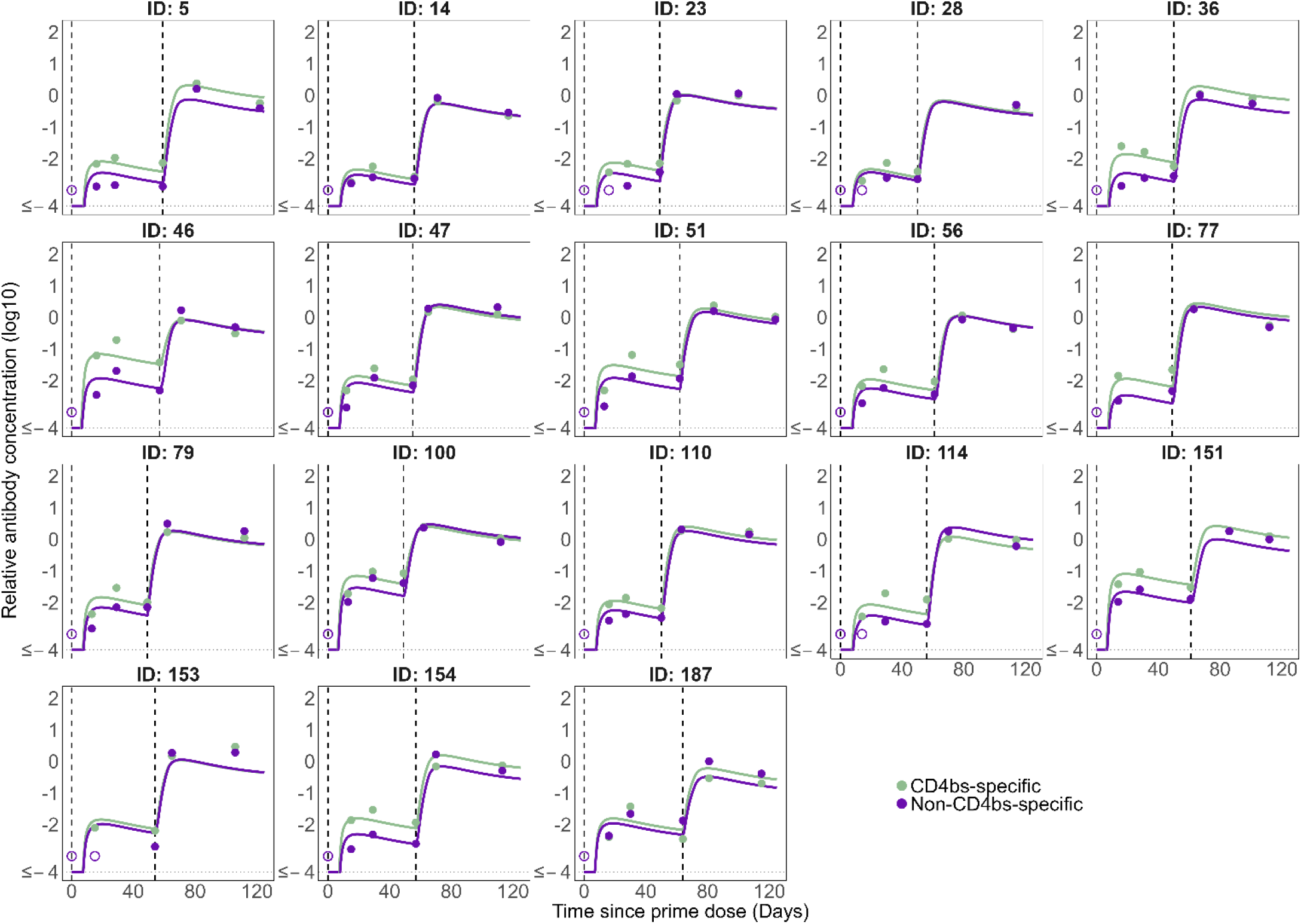
Individual-level predictions for antibody levels in the low-dose group. Observed (dots) and model-predicted (solid lines) antibody levels (log_10_) for participants in the low-dose group of the G001 trial. Results are shown for two antibody specificities: CD4bs (purple) and non-CD4bs (green). Vertical dashed lines indicate vaccinations.

**Figure SM3:**
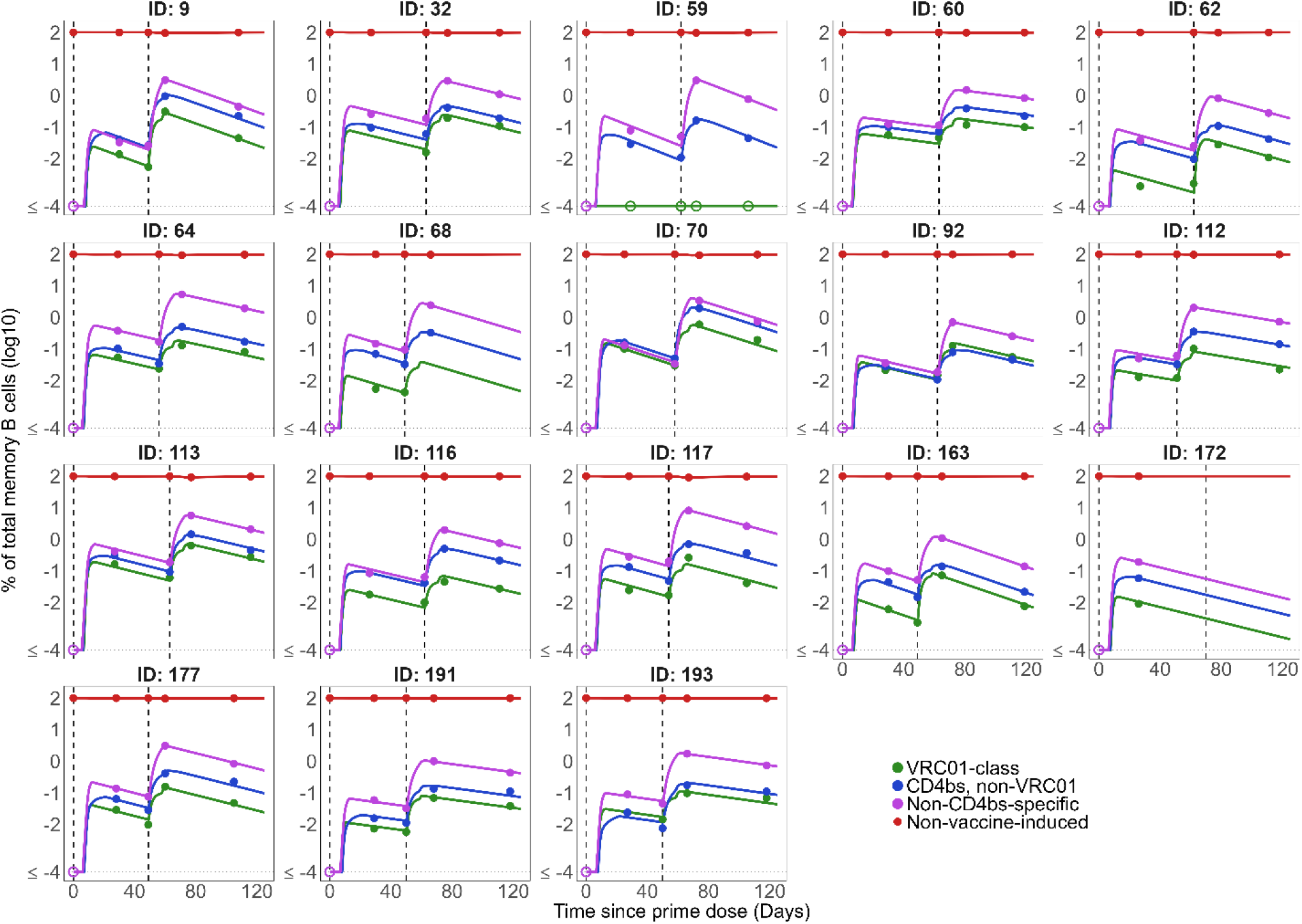
Individual-level predictions for memory B-cell frequencies in the high-dose group. Observed (dots) and model-predicted (solid lines) frequencies of four memory B-cell classes (log_10_) as a fraction of total memory B cells, for participants in the high-dose group of the G001 trial. The four classes are: VRC01-class (green), non-VRC01-class CD4bs (blue), non-CD4bs (purple), and non-vaccine-induced (red). Vertical dashed lines indicate vaccinations.

**Figure SM4:**
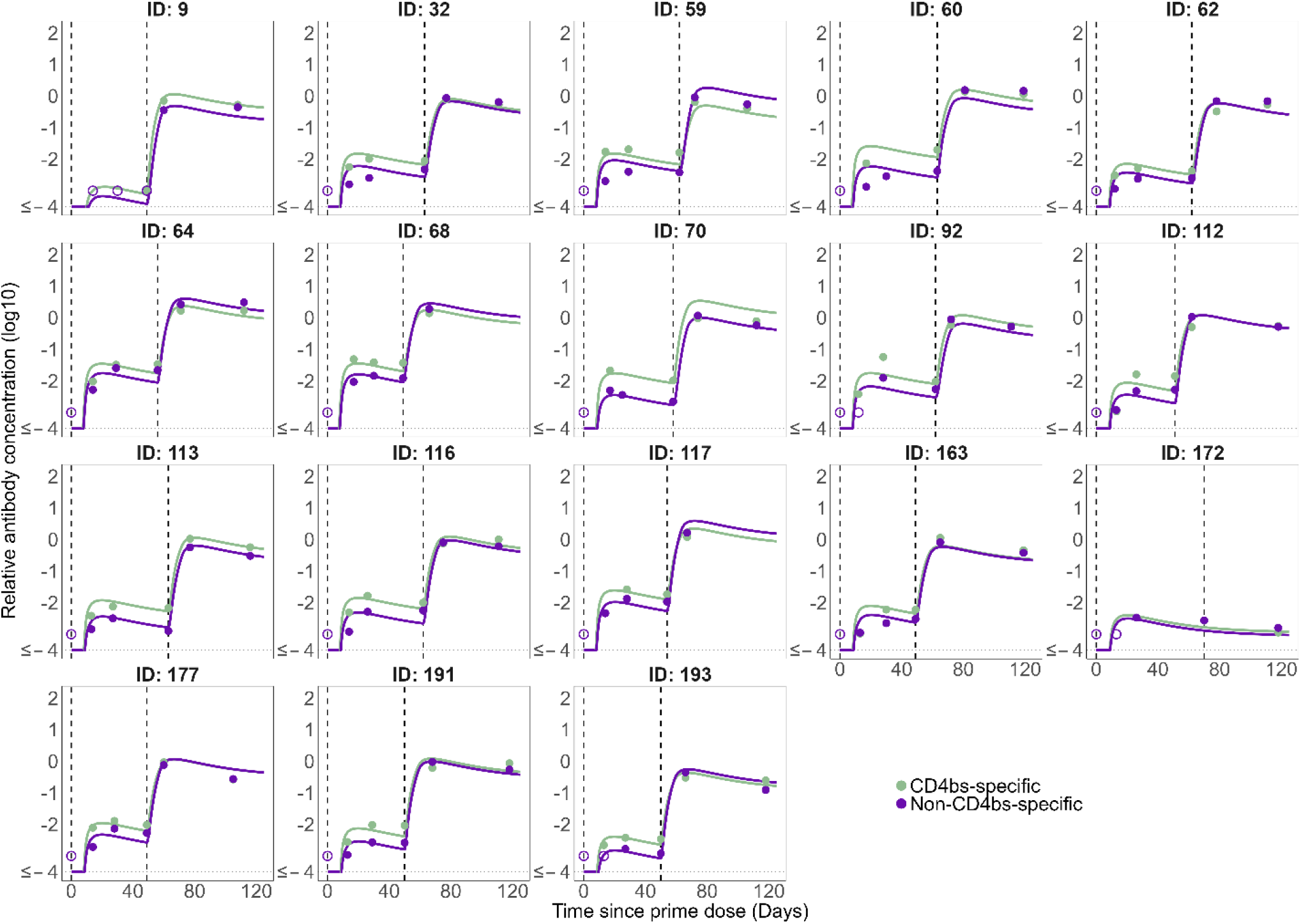
Individual-level predictions for antibody levels in the high-dose group. Observed (dots) and model-predicted (solid lines) antibody levels (log_10_) for participants in the high-dose group of the G001 trial. Results are shown for two antibody specificities: CD4bs (gold) and non-CD4bs (green). Vertical dashed lines indicate vaccinations.

**Table SM1:**
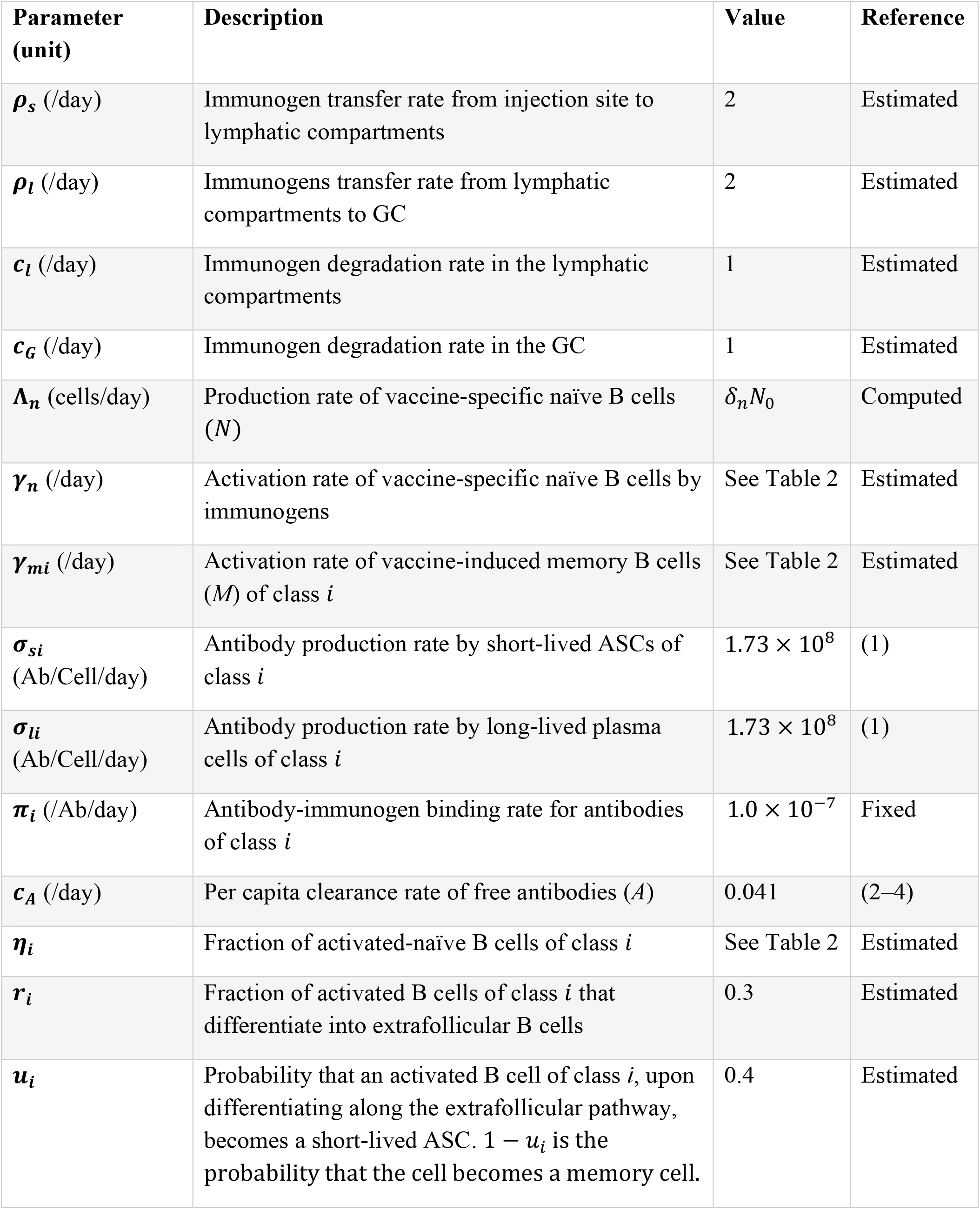
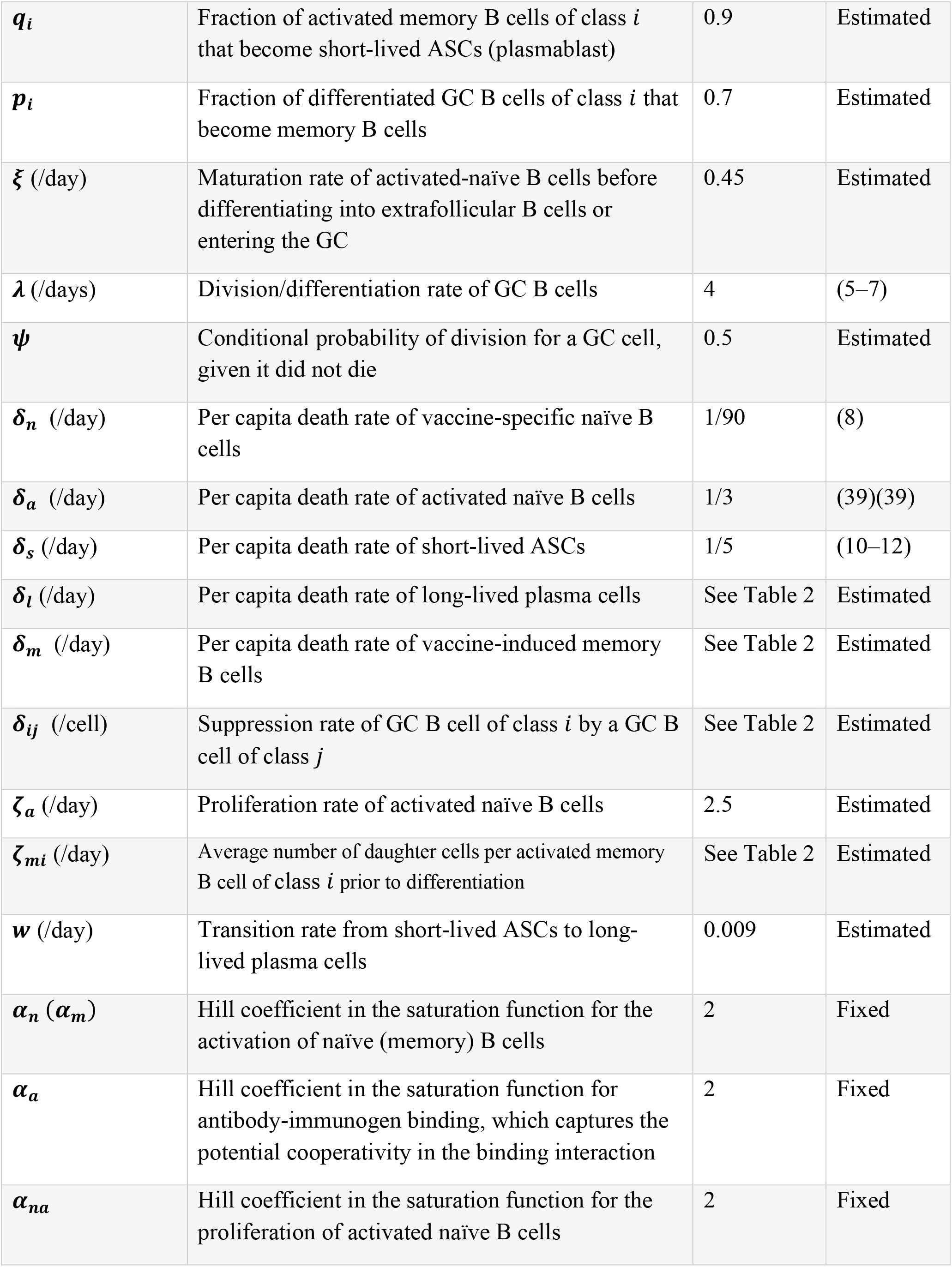
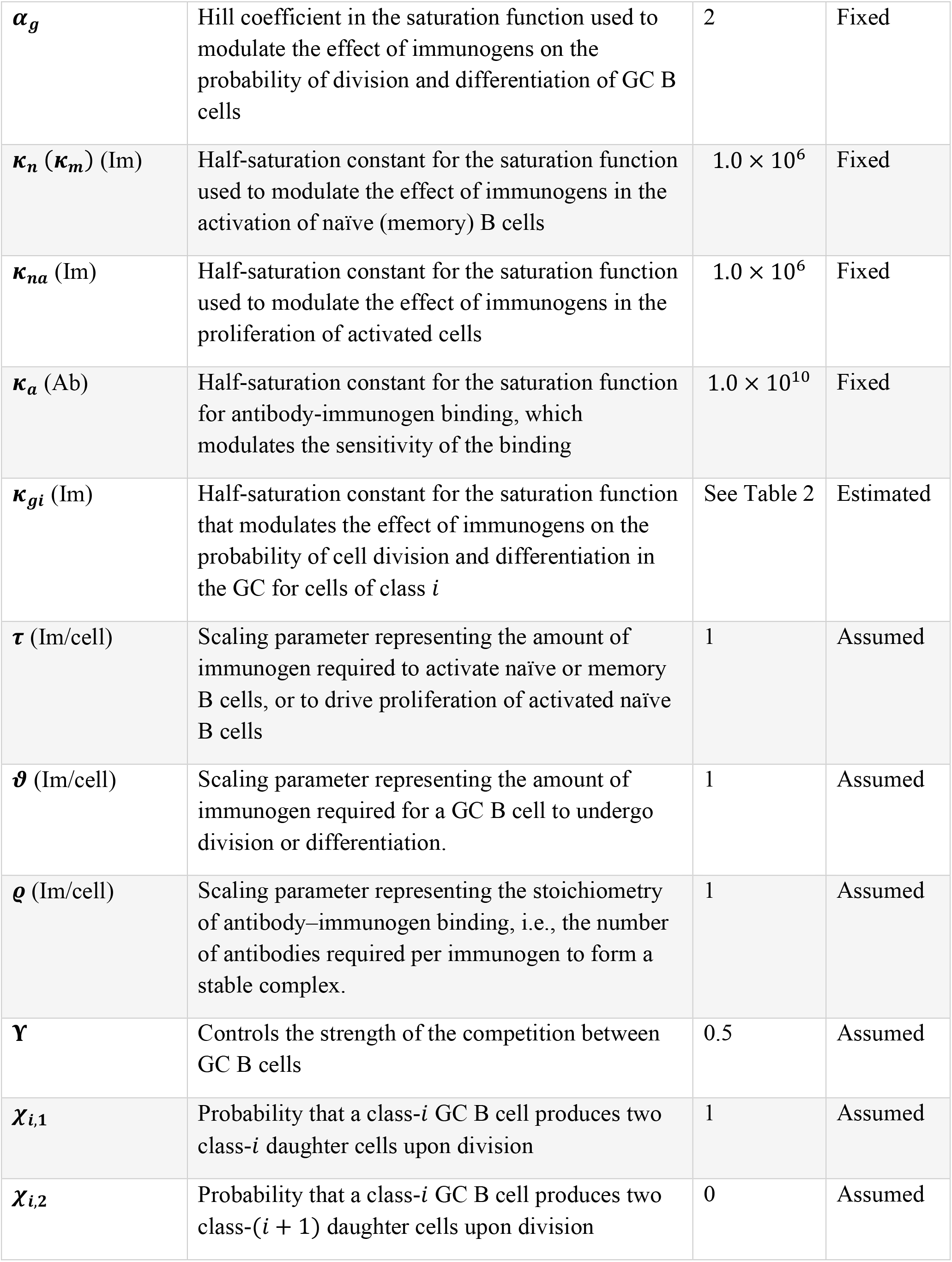
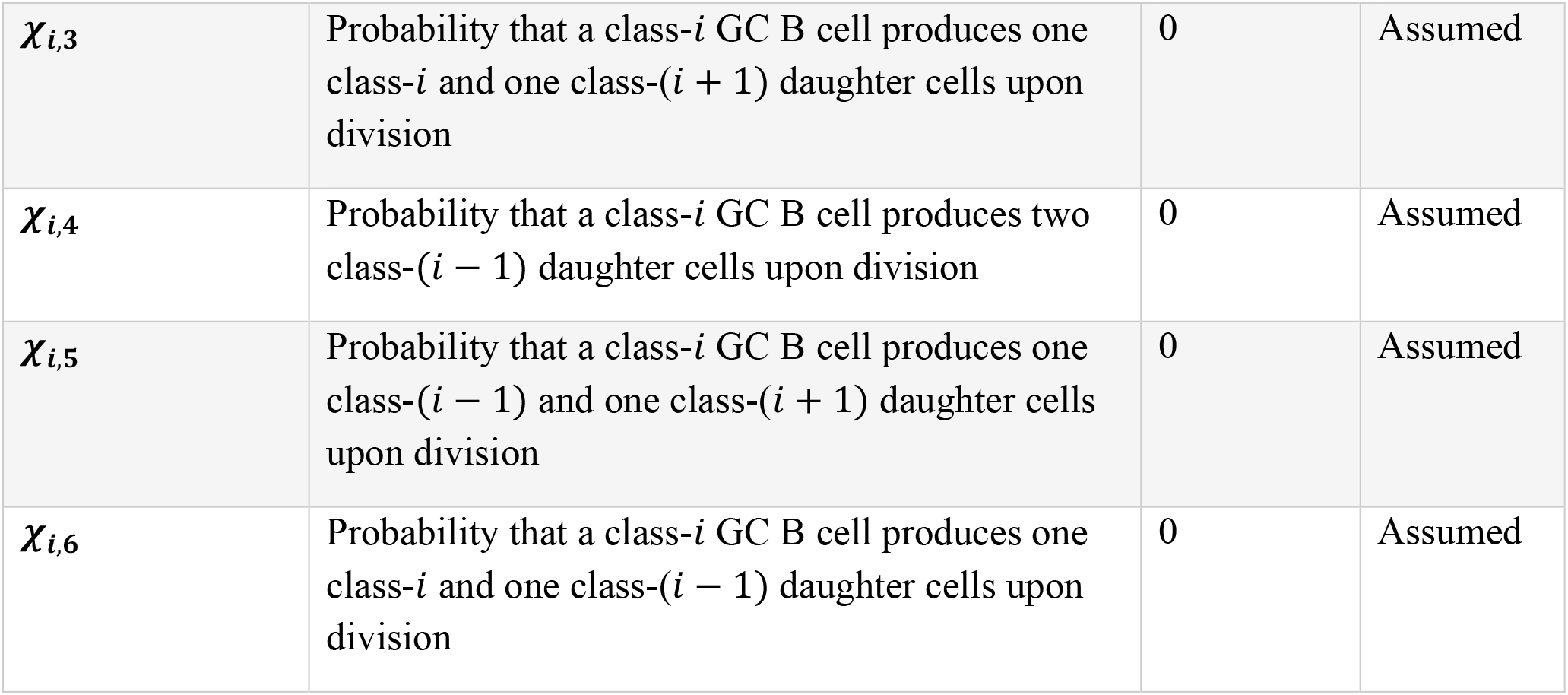
Model parameters and description.

| Parameter (unit) | Description | Value | Reference |
| --- | --- | --- | --- |
| $\rho_s$ (/day) | Immunogen transfer rate from injection site to lymphatic compartments | 2 | Estimated |
| $\rho_l$ (/day) | Immunogens transfer rate from lymphatic compartments to GC | 2 | Estimated |
| $c_l$ (/day) | Immunogen degradation rate in the lymphatic compartments | 1 | Estimated |
| $c_G$ (/day) | Immunogen degradation rate in the GC | 1 | Estimated |
| $\Lambda_n$ (cells/day) | Production rate of vaccine-specific naïve B cells ($N$) | $\delta_n N_0$ | Computed |
| $\gamma_n$ (/day) | Activation rate of vaccine-specific naïve B cells by immunogens | See Table 2 | Estimated |
| $\gamma_{mi}$ (/day) | Activation rate of vaccine-induced memory B cells ($M$) of class $i$ | See Table 2 | Estimated |
| $\sigma_{si}$ (Ab/Cell/day) | Antibody production rate by short-lived ASCs of class $i$ | $1.73 \times 10^8$ | (1) |
| $\sigma_{li}$ (Ab/Cell/day) | Antibody production rate by long-lived plasma cells of class $i$ | $1.73 \times 10^8$ | (1) |
| $\pi_i$ (/Ab/day) | Antibody-immunogen binding rate for antibodies of class $i$ | $1.0 \times 10^{-7}$ | Fixed |
| $c_A$ (/day) | Per capita clearance rate of free antibodies ($A$) | 0.041 | (2–4) |
| $\eta_i$ | Fraction of activated-naïve B cells of class $i$ | See Table 2 | Estimated |
| $r_i$ | Fraction of activated B cells of class $i$ that differentiate into extrafollicular B cells | 0.3 | Estimated |
| $u_i$ | Probability that an activated B cell of class $i$, upon differentiating along the extrafollicular pathway, becomes a short-lived ASC. $1 - u_i$ is the probability that the cell becomes a memory cell. | 0.4 | Estimated |
| $q_i$ | Fraction of activated memory B cells of class $i$ that become short-lived ASCs (plasmablast) | 0.9 | Estimated |
| $p_i$ | Fraction of differentiated GC B cells of class $i$ that become memory B cells | 0.7 | Estimated |
| $\xi$ (/day) | Maturation rate of activated-naïve B cells before differentiating into extrafollicular B cells or entering the GC | 0.45 | Estimated |
| $\lambda$ (/days) | Division/differentiation rate of GC B cells | 4 | (5–7) |
| $\psi$ | Conditional probability of division for a GC cell, given it did not die | 0.5 | Estimated |
| $\delta_n$ (/day) | Per capita death rate of vaccine-specific naïve B cells | 1/90 | (8) |
| $\delta_a$ (/day) | Per capita death rate of activated naïve B cells | 1/3 | (39)(39) |
| $\delta_s$ (/day) | Per capita death rate of short-lived ASCs | 1/5 | (10–12) |
| $\delta_l$ (/day) | Per capita death rate of long-lived plasma cells | See Table 2 | Estimated |
| $\delta_m$ (/day) | Per capita death rate of vaccine-induced memory B cells | See Table 2 | Estimated |
| $\delta_{ij}$ (/cell) | Suppression rate of GC B cell of class $i$ by a GC B cell of class $j$ | See Table 2 | Estimated |
| $\zeta_a$ (/day) | Proliferation rate of activated naïve B cells | 2.5 | Estimated |
| $\zeta_{mi}$ (/day) | Average number of daughter cells per activated memory B cell of class $i$ prior to differentiation | See Table 2 | Estimated |
| $w$ (/day) | Transition rate from short-lived ASCs to long-lived plasma cells | 0.009 | Estimated |
| $\alpha_n$ ($\alpha_m$) | Hill coefficient in the saturation function for the activation of naïve (memory) B cells | 2 | Fixed |
| $\alpha_a$ | Hill coefficient in the saturation function for antibody-immunogen binding, which captures the potential cooperativity in the binding interaction | 2 | Fixed |
| $\alpha_{na}$ | Hill coefficient in the saturation function for the proliferation of activated naïve B cells | 2 | Fixed |
| $\alpha_g$ | Hill coefficient in the saturation function used to modulate the effect of immunogens on the probability of division and differentiation of GC B cells | 2 | Fixed |
| $\kappa_n (\kappa_m)$ (Im) | Half-saturation constant for the saturation function used to modulate the effect of immunogens in the activation of naïve (memory) B cells | $1.0 \times 10^6$ | Fixed |
| $\kappa_{na}$ (Im) | Half-saturation constant for the saturation function used to modulate the effect of immunogens in the proliferation of activated cells | $1.0 \times 10^6$ | Fixed |
| $\kappa_a$ (Ab) | Half-saturation constant for the saturation function for antibody-immunogen binding, which modulates the sensitivity of the binding | $1.0 \times 10^{10}$ | Fixed |
| $\kappa_{gi}$ (Im) | Half-saturation constant for the saturation function that modulates the effect of immunogens on the probability of cell division and differentiation in the GC for cells of class $i$ | See Table 2 | Estimated |
| $\tau$ (Im/cell) | Scaling parameter representing the amount of immunogen required to activate naïve or memory B cells, or to drive proliferation of activated naïve B cells | 1 | Assumed |
| $\vartheta$ (Im/cell) | Scaling parameter representing the amount of immunogen required for a GC B cell to undergo division or differentiation. | 1 | Assumed |
| $\varrho$ (Im/cell) | Scaling parameter representing the stoichiometry of antibody-immunogen binding, i.e., the number of antibodies required per immunogen to form a stable complex. | 1 | Assumed |
| $\Upsilon$ | Controls the strength of the competition between GC B cells | 0.5 | Assumed |
| $\chi_{i,1}$ | Probability that a class-$i$ GC B cell produces two class-$i$ daughter cells upon division | 1 | Assumed |
| $\chi_{i,2}$ | Probability that a class-$i$ GC B cell produces two class-$(i + 1)$ daughter cells upon division | 0 | Assumed |
| $\chi_{i,3}$ | Probability that a class-$i$ GC B cell produces one class-$i$ and one class-$(i + 1)$ daughter cells upon division | 0 | Assumed |
| $\chi_{i,4}$ | Probability that a class-$i$ GC B cell produces two class-$(i - 1)$ daughter cells upon division | 0 | Assumed |
| $\chi_{i,5}$ | Probability that a class-$i$ GC B cell produces one class-$(i - 1)$ and one class-$(i + 1)$ daughter cells upon division | 0 | Assumed |
| $\chi_{i,6}$ | Probability that a class-$i$ GC B cell produces one class-$i$ and one class-$(i - 1)$ daughter cells upon division | 0 | Assumed |

### SM5. Establishing the structural identifiability of GC B-cell pairwise competition coefficients

This section proves Theorem 1, which establishes the structural identifiability of the pairwise competition coefficients *δ*_*ij*_, *i, j* = 1, …,3, between GC B-cell classes, when all other model parameters are fixed. The proof proceeds in two stages. The first stage reduces the uniqueness of the parameters *δ* = *δ*_*ij*_, *i, j* = 1, …,3 to a linear-independence property of the functions *G*_1_, *G*_2_, *G*_3_. Specifically, by (S6), if the G001 Models parametrized by *δ* and *δ*′ = *δ*′_*ij*_, *i, j* = 1, …,3 produce the same solution, then, for each *i*,

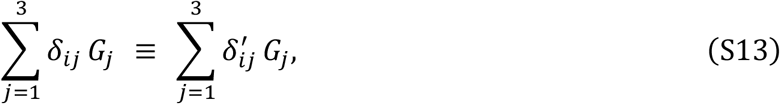

on a neighborhood of 0^+^. If the functions *G*_1_, *G*_2_, *G*_3_ are linearly independent on this neighborhood, Eq. (S13) implies *δ*_*ij*_ = δ^′^ for *j* = 1, 2, 3, and for every *i*, and hence *δ* = *δ*′. The second stage establishes this linear independence. We derive the local expansion of *G*_1_, *G*_2_, *G*_3_ near *t* = 0 and show that no nontrivial linear combination of these expansions can vanish. This is the content of Theorem 2, which we prove first. Once Theorem 2 is established, the linear independence of *G*_1_, *G*_2_, *G*_3_, together with (S13), yields *δ* = *δ*′. This completes the first stage of the argument and, consequently, the proof of Theorem 1. The remainder of this section is devoted first to Theorem 2 and then to returning to Theorem 1 at the end.

#### Linear independence of *G*_1_(*t*), *G*_2_(*t*), and *G*_3_(*t*) near *t* = 0

Let *t* ∈ [0, *T*] for some *T* ∈ (0, *t*_2_), where *t*_2_ denotes the time at which the second vaccination occurred. For any *κ, α* > 0, let *H*_*κ,α*_(*x*) : = *x*^*α*^/(*x*^*α*^ + *κ*^*α*^), and write 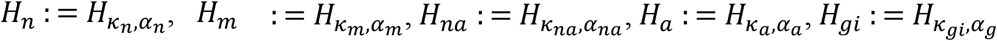, we have that *I*_*S*_(0) = *I*_*S*1_ > 0, *N*(0) = *N*_0_ : = *Λ*_*n*_/*δ*_*n*_, and *I*_*L*_, *I*_*G*_, *N*_*a*_, *G*_*i*_, *P*_*si*_, *P*_*li*_, *M*_*i*_, *A*_*i*_ are all 0. All rate constants, half-saturation constants, and Hill exponents are positive; *η*_*i*_, *r*_*i*_, *q*_*i*_, *u*_*i*_, *p*_*i*_, *φ*_*i*_, *ψ*_*i*_ ∈ [0,1] with *η*_*i*_(1 − *r*_*i*_) > 0; δ_*ij*_ ≥ 0 and ϒ > 0. We assume that a continuous, nonnegative solution of (S1)-(S11) exists on [0, *T*].

Define *a* : = *ρ*_*s*1_ *I*_*S*1_, *µ* : = min{1, *α*_*n*_},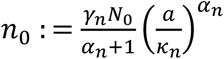, and

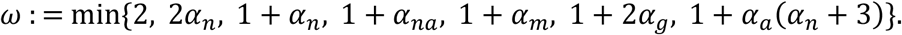

In this analysis, the notations *O*(⋅) and *o*(⋅) refer to the limit *t* → 0^+^, with constants depending only on the model parameters and on *T*.

##### Theorem 2.

*Assume* min (*α*_*m*_, 2*α*_*g*_)> 1. *For k*= 0, 1, 2 *and t*> 0 *put* 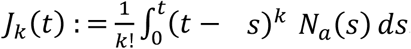. *Then, for i* = 1, 2, 3,

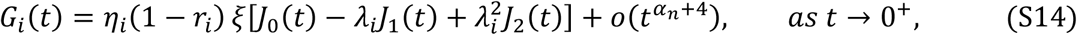

*where*

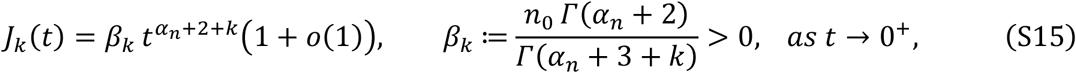

*with* 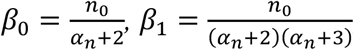, *and* 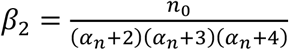. *Consequently, if λ*_1_, *λ*_2,_ *λ*_3_ *are pairwise distinct, then G*_1_, *G*_2_, *G*_3_ *are linearly independent on every interval* (0, *T*).

**Proof of Theorem 2**

In this proof, we use four elementary facts, stated below.

**F1**. Let *X*: = (*x*/*κ*)^*α*^. Then *H*_*κ,α*_(*xx*) = *X* − *X*^2^/(1 + *X*) and, when *x* ≥ 0, 0 ≤ *X* − *H*_*κ,α*_(*x*) ≤ *X*^2^; in particular *H*_*κ,α*_(*x*) = (*x*/*κ*)^*α*^ + *O*((*x*/*κ*)^2*α*^) and *H*_*κ,α*_(*x*) ≤ min(*X*, 1).

**F2**. (1 + *ε*)^*α*^ = 1 + *αε* + *O*(*ε*^2^) for 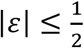.

**F3**. For an integer *k* ≥ 0 and *β* > −1, substituting *s*= *tx*,

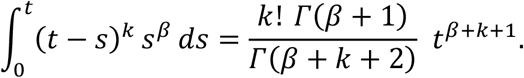

**F4**. For 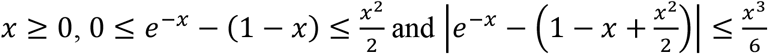.

#### Step 1: preliminary bounds

We first develop bounds describing the behavior near *t*= 0 of certain functions appearing in the ODEs that define the G001 Model; these bounds will allow us to characterize the behavior of *G*_*i*_(*t*) near *t* = 0. To shorten notation, we use *f*′ to denote the first derivative of any univariate function *f*. First, at any time *t* ∈ [0, *T*], only the first immunization dose has been administered. Hence, by (S1), 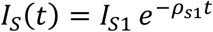. Dropping the nonpositive terms in (S2) and (S3) gives *I*_*L*_′ ≤ *ρ*_*s*1_*I*_*S*_ ≤ *a* and *I*_*G*_′ ≤ *ρ* _*l*_ *I*_*L*_, so that *I*_*L*_ ≤ *at* and 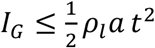. Then (S5) and F1 give 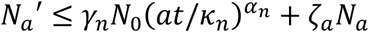. Integrating this inequality and using the fact that *N*_*a*_(0) = 0, we obtain that 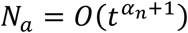. In (S6), (S8) and (S10), all coefficients are bounded (*H* ≤ 1, *Φ*_*i*_≤ 1), so the derivative of ∑_*i*_ (*G*_*i*_ + *M*_*i*_ + *P*_*si*_) is at most *c*_0_ (*N*_*a*_ + ∑_*i*_ (*G*_*i*_ + *M*_*i*_ + *P*_*si*_)) for some constant *c*_0_ > 0. Proceeding as done with *N*_*a*_, we obtain that *G*_*i*_, *M*_*i*_, 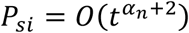, and then (S9) and (S11) give 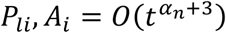. Therefore, every nonpositive term in (S2) is *O*(*t*^*μ*^), and *ρ*_*s*1_*I*_*S*_ = *a* + *O*(*t*), such that *I*_*L*_′ = *a* + *O*(*t*^*μ*^); hence *I*_*L*_ = *at* (1 + *O*(*t*^*μ*^). Finally, (S4) gives 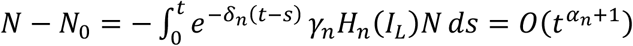,so *N*= *N*_*o*_( 1 + *o*(1)).

Next, integrating (S2) from 0 to *t* and approximating each term with the above bounds gives

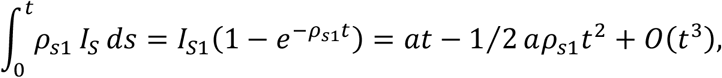

and

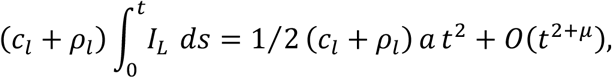

for the first and part of the second term, respectively. By F1 and F2, 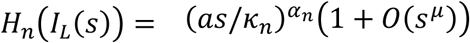. Using 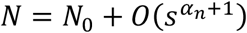, the naïve B cell activation term gives

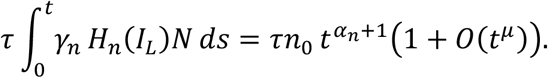

Finally, the antibody, memory B cell, and activated B cell terms can be shown to be 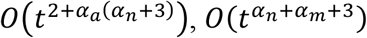 and 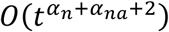, respectively. After dividing by *at*, every term is *O*(*t*^*ω*^). Hence,

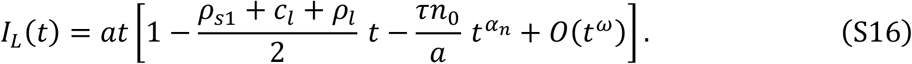

#### Step 2: Expansion of *N*_*a*_ near 0

Let *Q* : = *γ*_*n*_*H*_*n*_(*I*_*L*_)*N*. By Step 1, 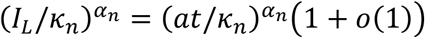, which tends to 0 as *t* → 0^+^. Hence, F1 gives 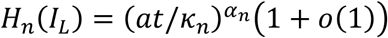. Since *N* = *N*_0_1 + *o*(1), we find that

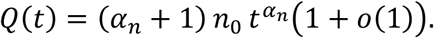

Using (S5) and *N*_*a*_(0) = 0,

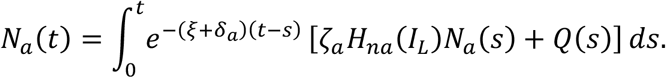

In the first term, 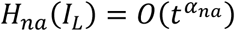 by F1 and 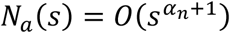 by Step 1. Therefore, 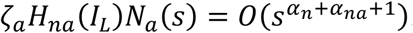 for *s* ∈ [0, *t*], integrating this term over [0, *t*] increases the order to 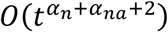, which is 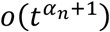.

For the second term, expand the kernel by one order. Put *c* : = *ξ* + *δ*_*a*_ > 0. By the first bound in F4 with *x*= *c*(*t* − *s*) ∈ [0, *ct*],

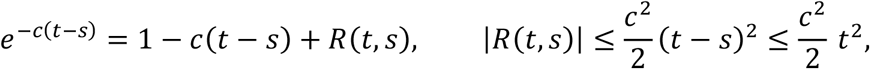

so that

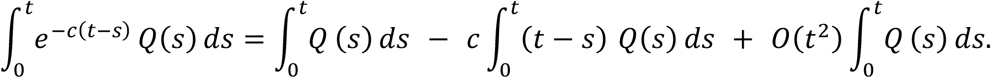

Write 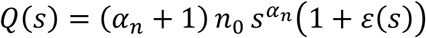 with *ε*(*s*) → 0 as *s* → 0^+^. By F3 with *β* = *α*_*n*_,

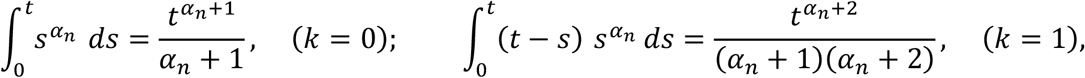

and in each case the *ε*-part is bounded by (sup_[0,*t*]_|*ε*|) times the same quantity. The three terms above are therefore

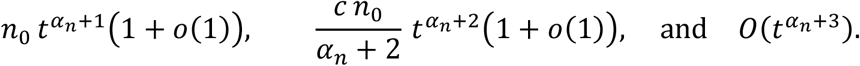

The last two are 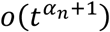, as is the ζ_*a*_ contribution. Hence, we can only retain the first, and

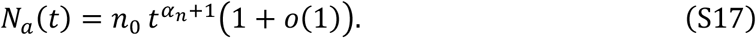

#### Step 3: Expansion of *G*_***m***_ near 0

Since *v*_*i*_ = *Ψ*_*i*_*Φ*_*i*_, (S6) can be expressed as

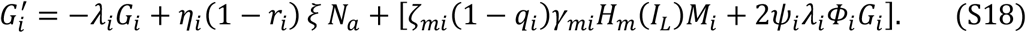

Using F1 and Step 1 gives 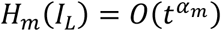 and 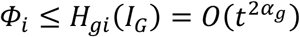, while 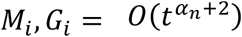 by Step 1. Hence, the order of the term in the bracket is 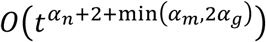. Since *G*_*i*_(0) = 0, integrating (S18) gives

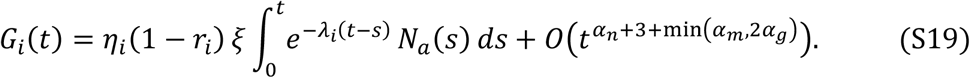

Using the Taylor expanding of 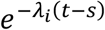 as t → 0^+^ in the integral in *G*_*i*_(*t*) together with 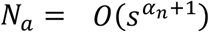, we obtain

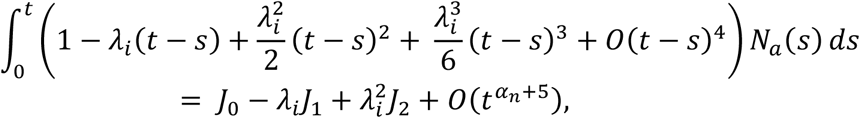

where 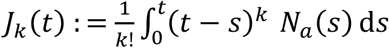. Substituting this result back into (S19), we obtain

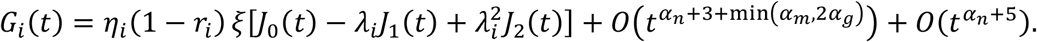

Because min (*α*_*m*_, 2*α*_*g*_)> 1, both error terms are 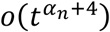, which gives *G*_*i*_(*t*) as presented in (S14). Finally, inserting (S17) into the definition of *J*_*k*_ and applying F3 with *β* = *α*_*n*_ + 1 gives (S15); the stated values of *β*_0_, *β*_1_, *β*_2_ follow from *Γ*(*z* + 1) = *zΓ*(*z*).

**Step 4: Linear independence of *G***_**1**_, ***G***_**2**_, **and *G***_**3**_

Assume that *λ*_1_, *λ*_2_, and *λ*_3_ are pairwise distinct, suppose ∑_*i*_ *c*_*i*_ *G*_*i*_ ≡ 0 on some (0, 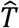), and set *x*_*i*_ : = *c*_*i*_ *η*_*i*_(1 − *r*_*i*_). Summing (S14) against the *c*_*i*_ yields

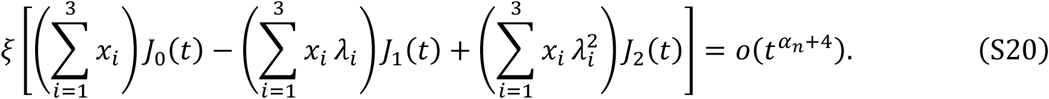

We extract the three sums in turn, using (S15). Divide (S20) by 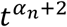 and let *t* → 0^+^. By (S15), 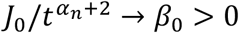, while 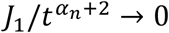 and 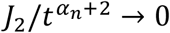, and the right-hand side tends to 0. Hence ∑_*i*_ *x*_*i*_ = 0. With ∑_*i*_ *x*_*i*_ = 0 the first term of (S20) vanishes identically, not merely to leading order. Divide what remains by 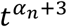 and let *t* → 0^+^: now 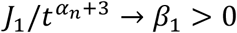 and 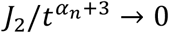, giving ∑_*i*_ *x*_*i*_ *λ*_*i*_ = 0. This makes the second term vanish. Dividing the remainder by 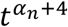 and letting *t* → 0^+^, with 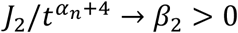, gives ∑_*i*_ *x*_*i*_ *λ*^2^ = 0. Thus, *x*_1_, *x*_2_, *x*_3_ solve the linear system

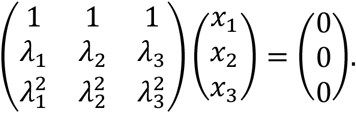

The determinant of the Vandermonde matrix is ∏_*i*<*j*_ *λ*_*j*_ − *λ*_*i*_ ≠ 0 because the *λ*_*i*_ are pairwise distinct. The system therefore has only the solution *x*_1_ = *x*_2_ = *x*_3_ = 0, and since *η*_*i*_(1 − *r*_*i*_) > 0 we get *c*_*i*_ = *x*_*i*_/*η*_*i*_(1 − *r*_*i*_) = 0 for every *i*. This completes the proof of Theorem 2.

**Proof of Theorem 1**

If *δ* = *δ*′ the two models coincide. Suppose now that the models parametrized by *δ* and *δ*′ produce the same solution of (S1)-(S11) on [0, *T*], and write *Φ*_*i*_ and *Φ*′_*i*_ for the two versions of the competition factor,

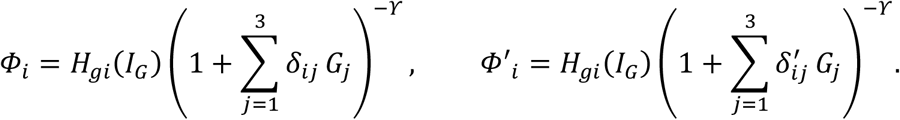

Fix *i*. Every term of (S6) other than 2*ψ*_*i*_*λ*_*i*_*Φ*_*i*_*G*_*i*_ is identical under the two parametrizations, as is 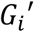; hence 2*ψ*_*i*_*λ*_*i*_*Φ*_*i*_*G*_*i*_ = 2*ψ*_*i*_*λ*_*i*_*Φ*′_*i*_*G*_*i*_ on [0, *T*]. Since *ψ*_*i*_ > 0 and *λ*_*i*_ > 0, this gives *Φ*_*i*_*G*_*i*_ = *Φ*′_*i*_*G*_*i*_.

By (S14) and (S15), 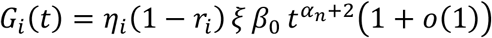 with *η*_*i*_(1 − *r*_*i*_) *ξ β*_0_ > 0, so there is *T*^*^ ∈ (0, *T*] with *G*_*i*_(*t*) > 0 for all *t* ∈ (0, *T*^*^). On that interval, we may cancel *G*_*i*_, so *Φ*_*i*_ = *Φ*′_*i*_. Moreover *I*_*G*_′ = *ρ*_*l*_*I*_*L*_ − *θ* ∑_*j*_ *λ*_*j*_ *v*_*j*_*G*_*j*_ − *c*_*G*_*I*_*G*_, and by Step 1 the first term is *ρ*_*l*_*at(*1 + *o*(1)) while the others are 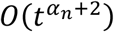 and *O*(*t*^2^); hence *I*_*G*_(*t*) > 0, and therefore *H*_*gi*_(*I*_*G*_) > 0, after shrinking *T*^*^ if necessary. Dividing by *H*_*gi*_(*I*_*G*_) leaves

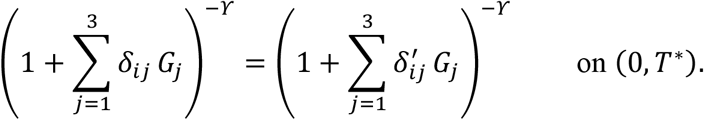

Both bases are at least 1 because 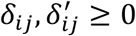 and *G*_*S*_ ≥ 0, and *u* ↦ *u*^−*ϒ*^ is strictly decreasing on (0, ∞) since *ϒ* > 0; the bases therefore coincide, which is (S13), that is,

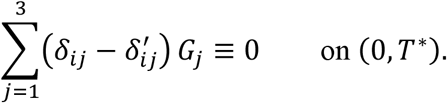

By Theorem 2, *G*_1_, *G*_2_, *G*_3_ are linearly independent on (0, *T*^*^), so 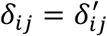 for *j* = 1, 2, 3. As *i* was arbitrary, *δ* = *δ*^′^. This completes the proof of Theorem 1. ∎

### SM6. Profile likelihood for the GC B-cell competition coefficients

**Figure SM5.**
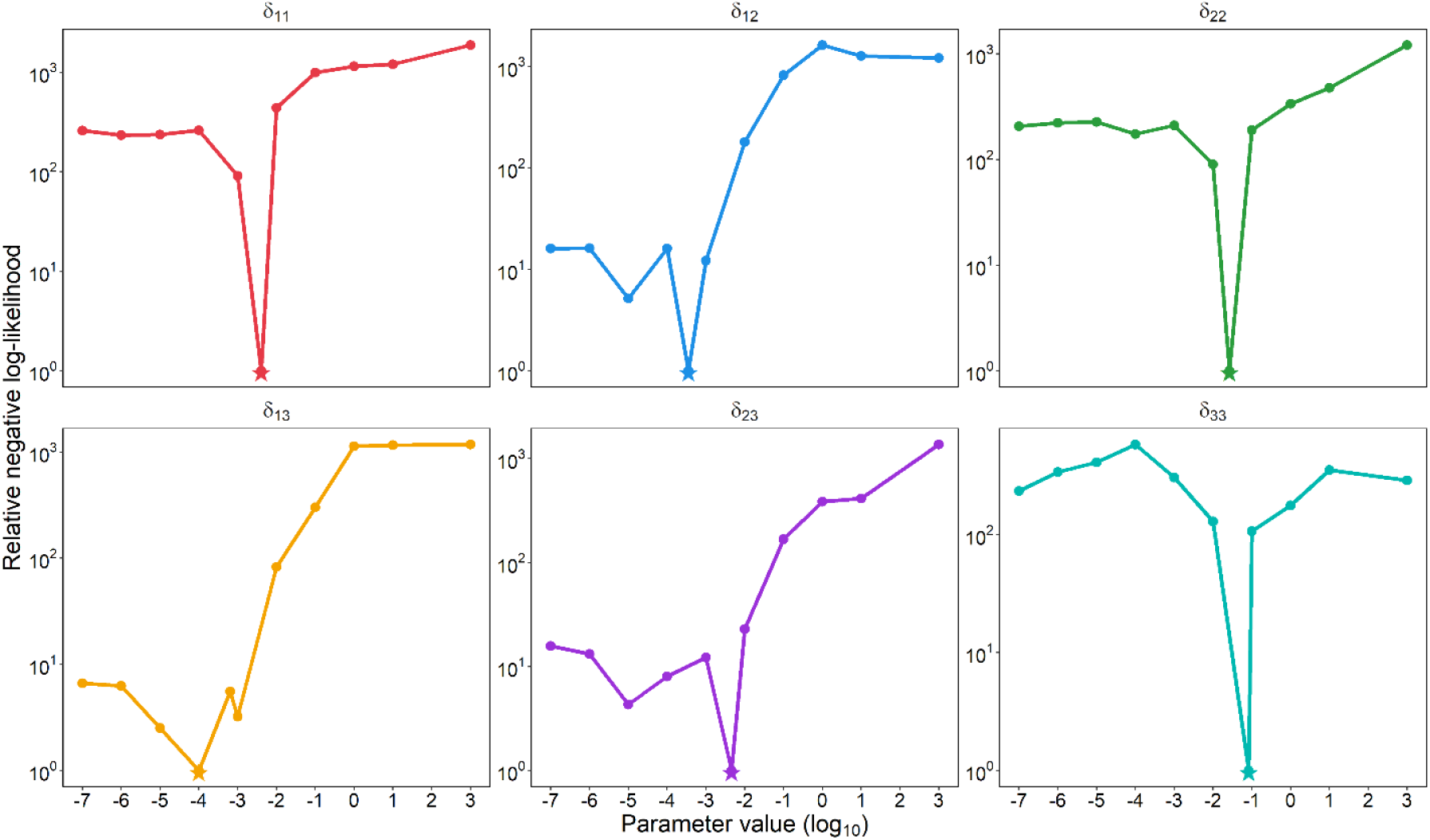
Profile-likelihood results for the germinal center B-cell competition coefficients. Each panel shows the negative log-likelihood (NLL) for each GC B-cell competition coefficient (log_10_ scale) at different values of the parameter, relative to the minimum NLL over the entire parameter set (offset by 1 for display on a logarithmic scale). The star marks the estimated value of each competition coefficient reported in Table 2.

**Figure SM6.**
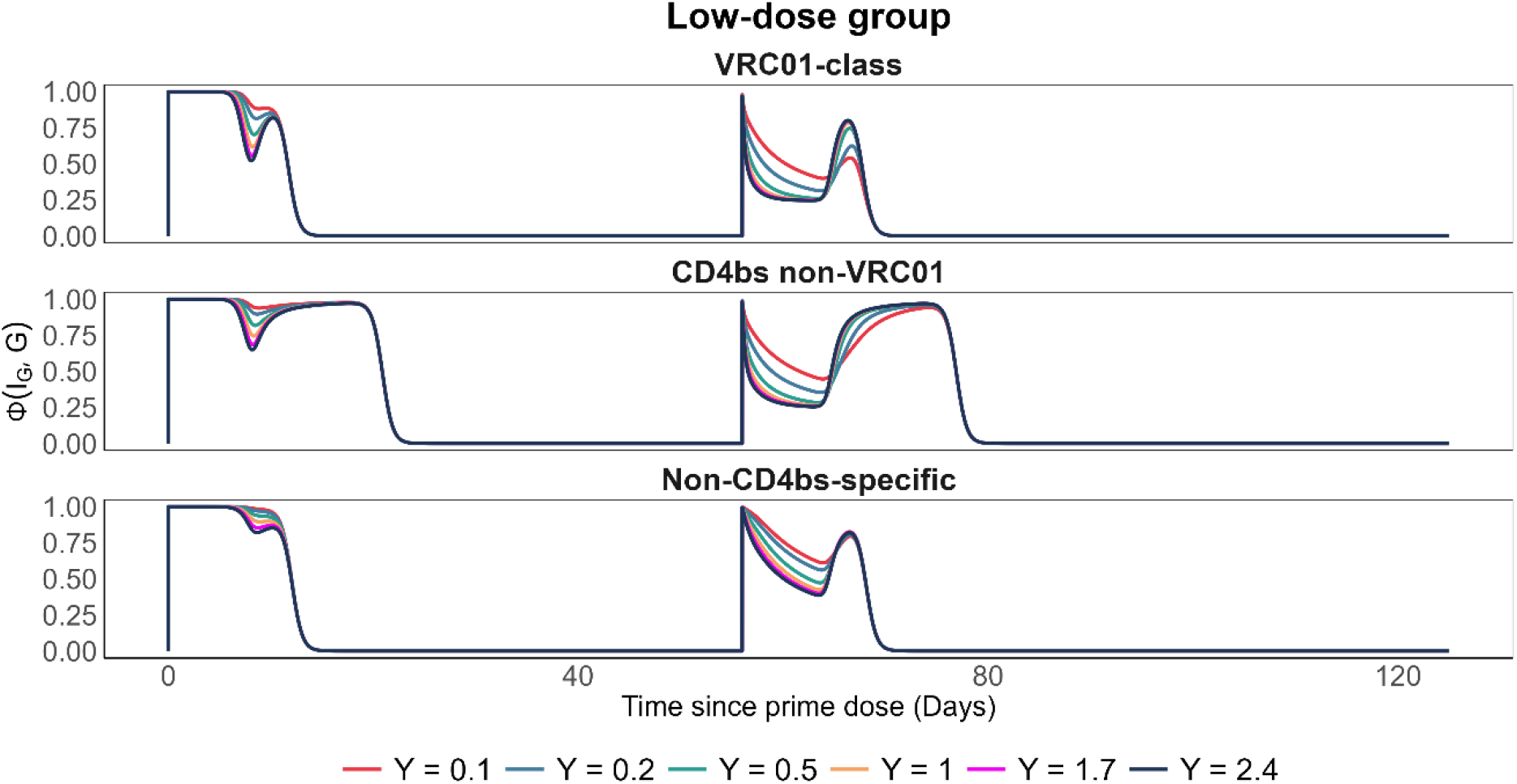
Simulated trajectories for the probability of division or differentiation (low-dose group). Probability of division or differentiation Φ(*I*_*G*_, *G*) over time (days) for the three B-cell classes (VRC01-class, CD4bs non-VRC01, and non-CD4bs-specific) in the low-dose group, shown for six representative values of ϒ.

**Figure SM7.**
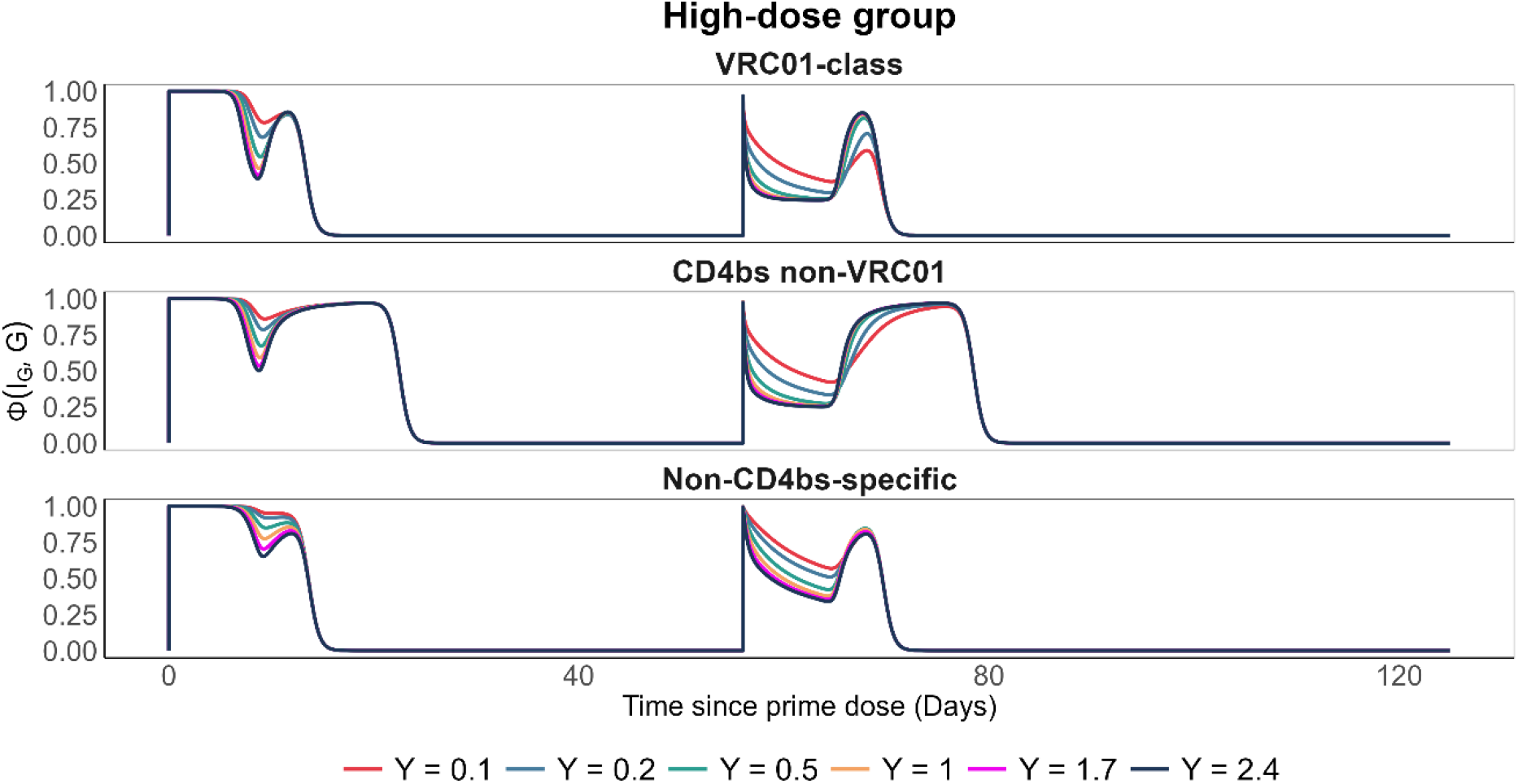
Simulated trajectories for the probability of division or differentiation (high-dose group). Probability of division or differentiation Φ(*I*_*G*_, *G*) over time (days) for the three B-cell classes (VRC01-class, CD4bs non-VRC01, and non-CD4bs-specific) in the low-dose group, shown for six representative values of ϒ.

**Figure SM8.**
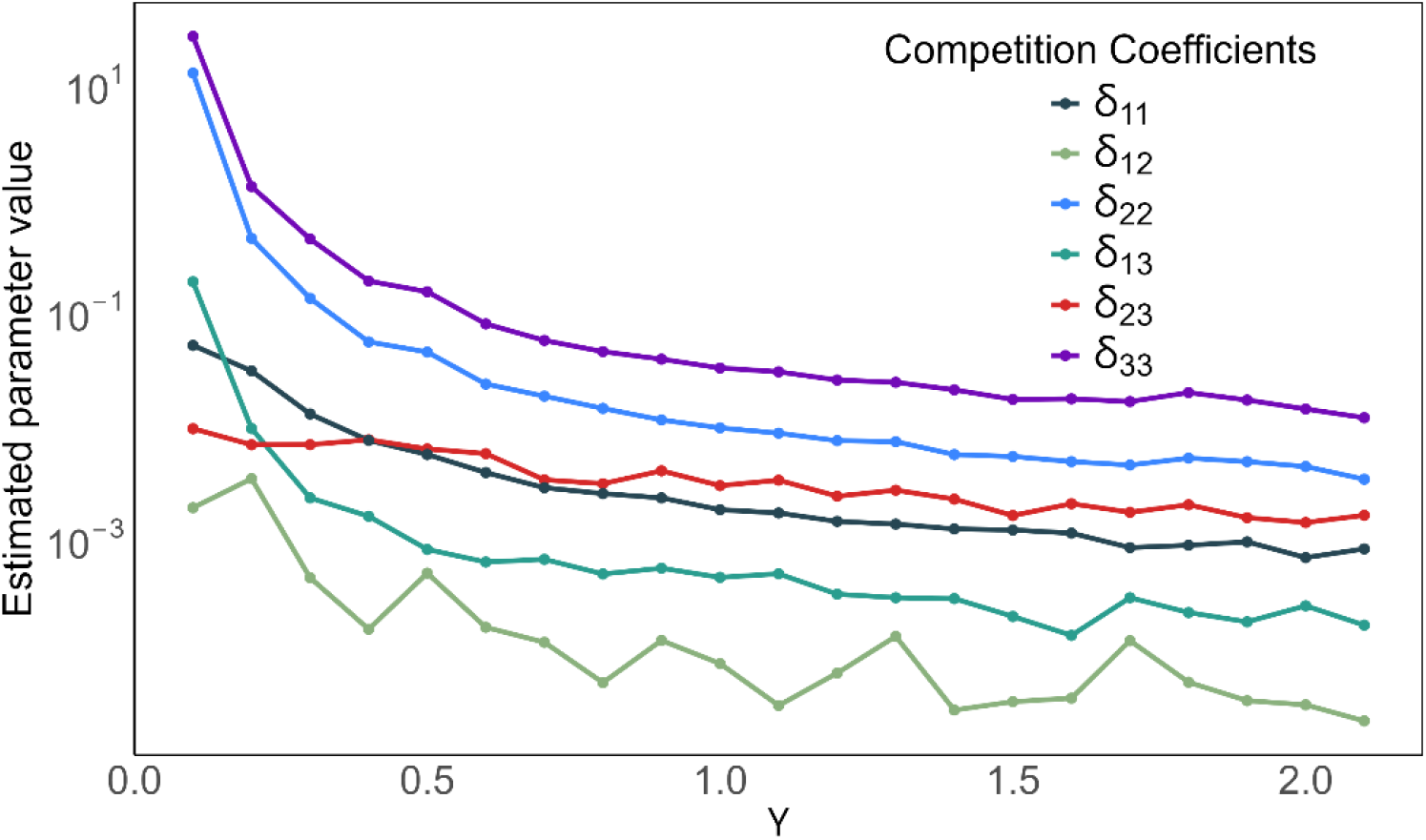
Estimated GC B cells competition coefficients as a function of ϒ.

### SM7. Effect of antibody masking on immune response dynamics

**Figure SM9:**
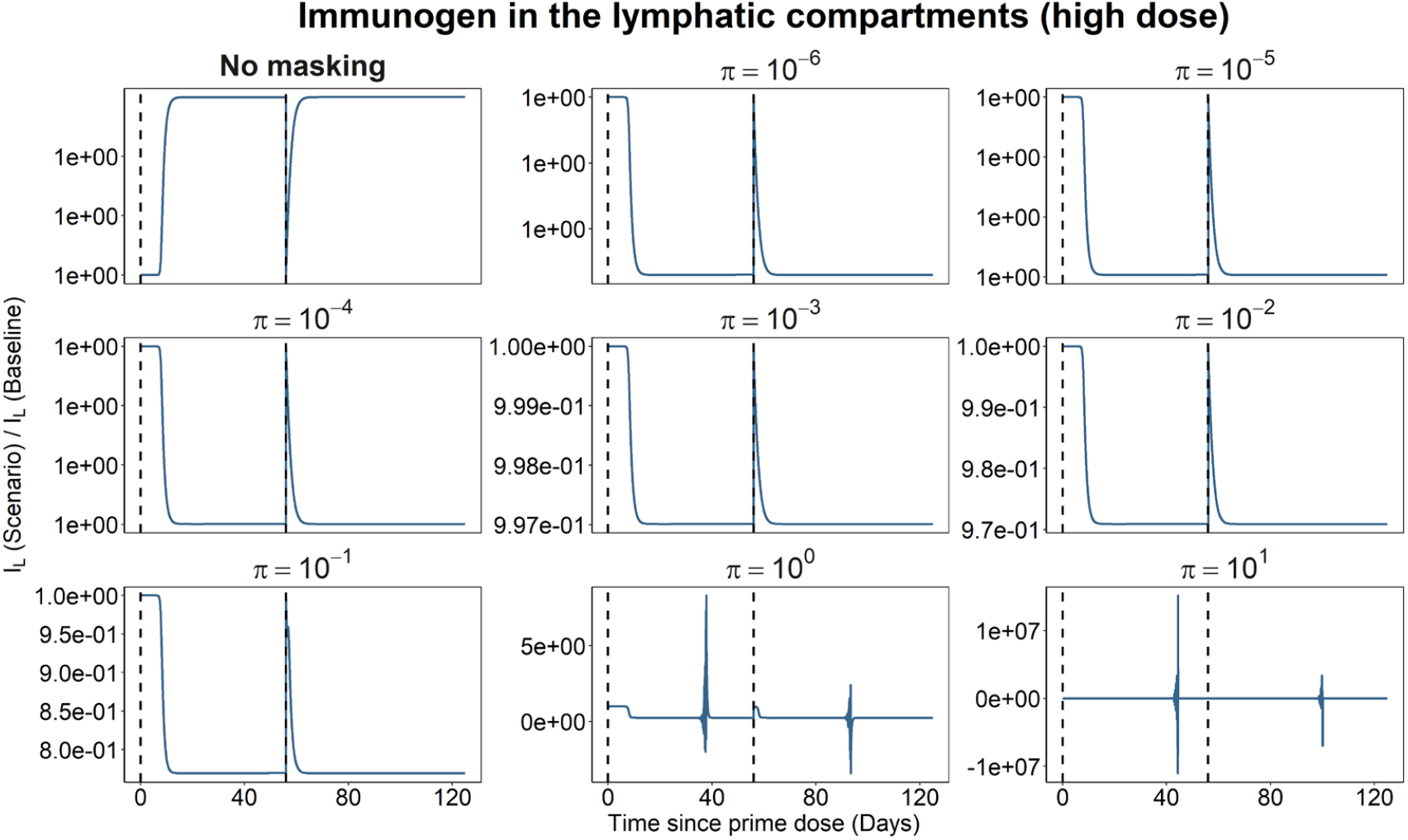
Effect of antibody masking on immunogen dynamics in the lymphatic compartment. Ratio of immunogen levels in the lymphatic compartment for different values of the immunogen–antibody binding rate (*π*) relative to the immunogen level at *π* = 10^−7^ (baseline).

**Figure SM10:**
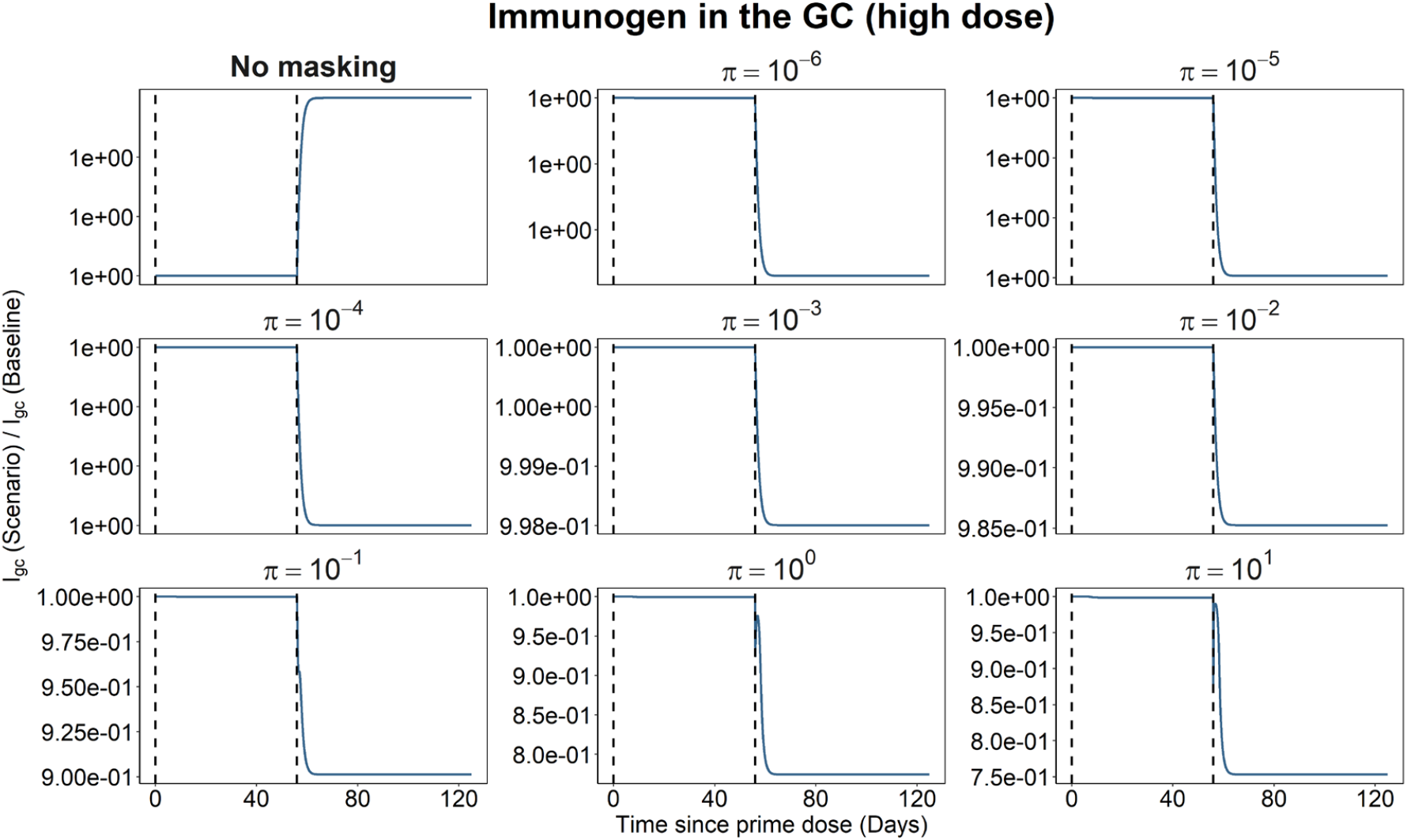
Effect of antibody masking on immunogen dynamics in the GC. Ratio of immunogen levels in the GC for different values of the immunogen–antibody binding rate (*π*) relative to the immunogen level at *π* = 10^−7^ (baseline).

**Figure SM11:**
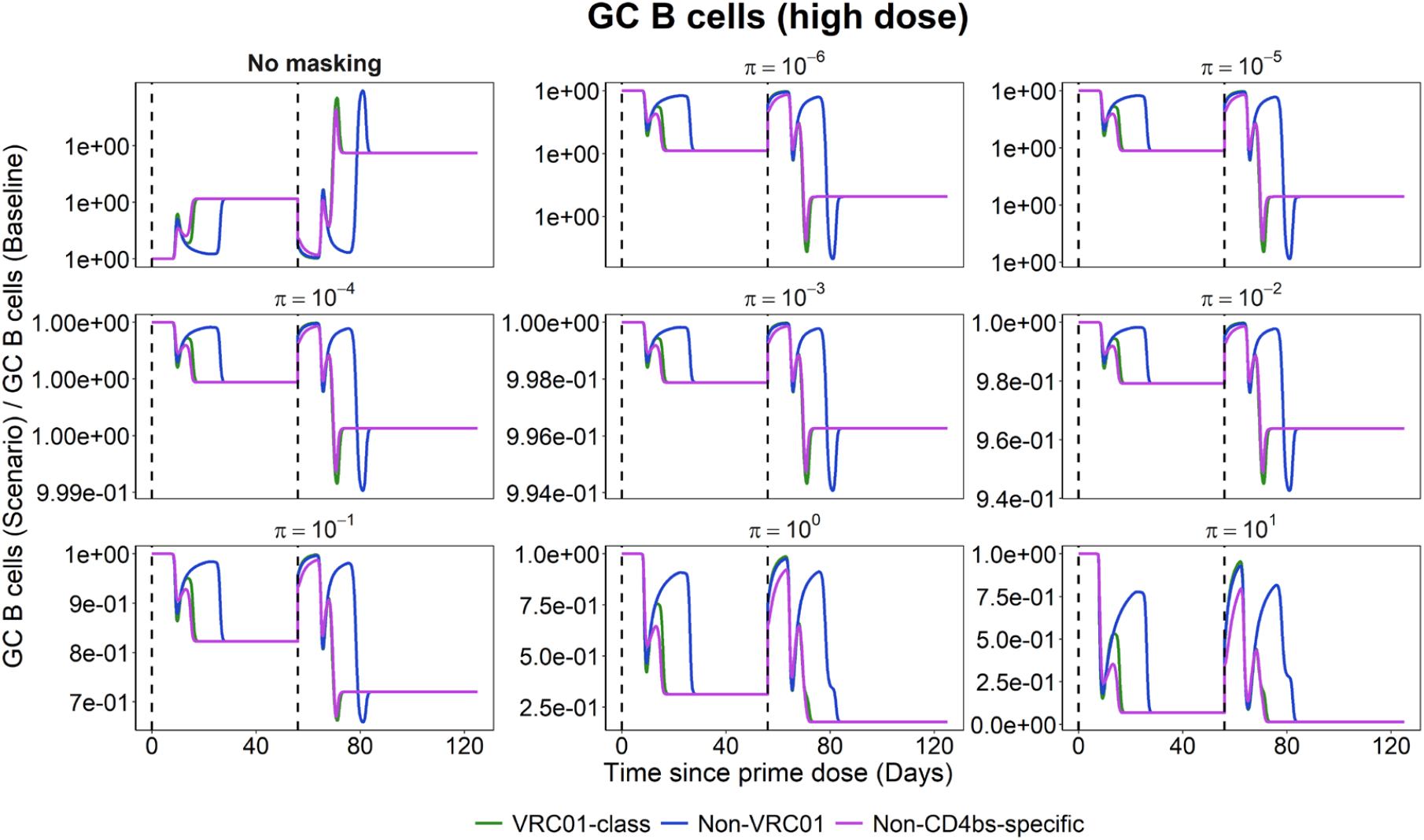
Effect of antibody masking on vaccine-induced GC B-cell dynamics. Ratio of GC B cells for different values of the immunogen–antibody binding rate (*π*) relative to the GC B cells at *π* = 10^−7^ (baseline).

**Figure SM12:**
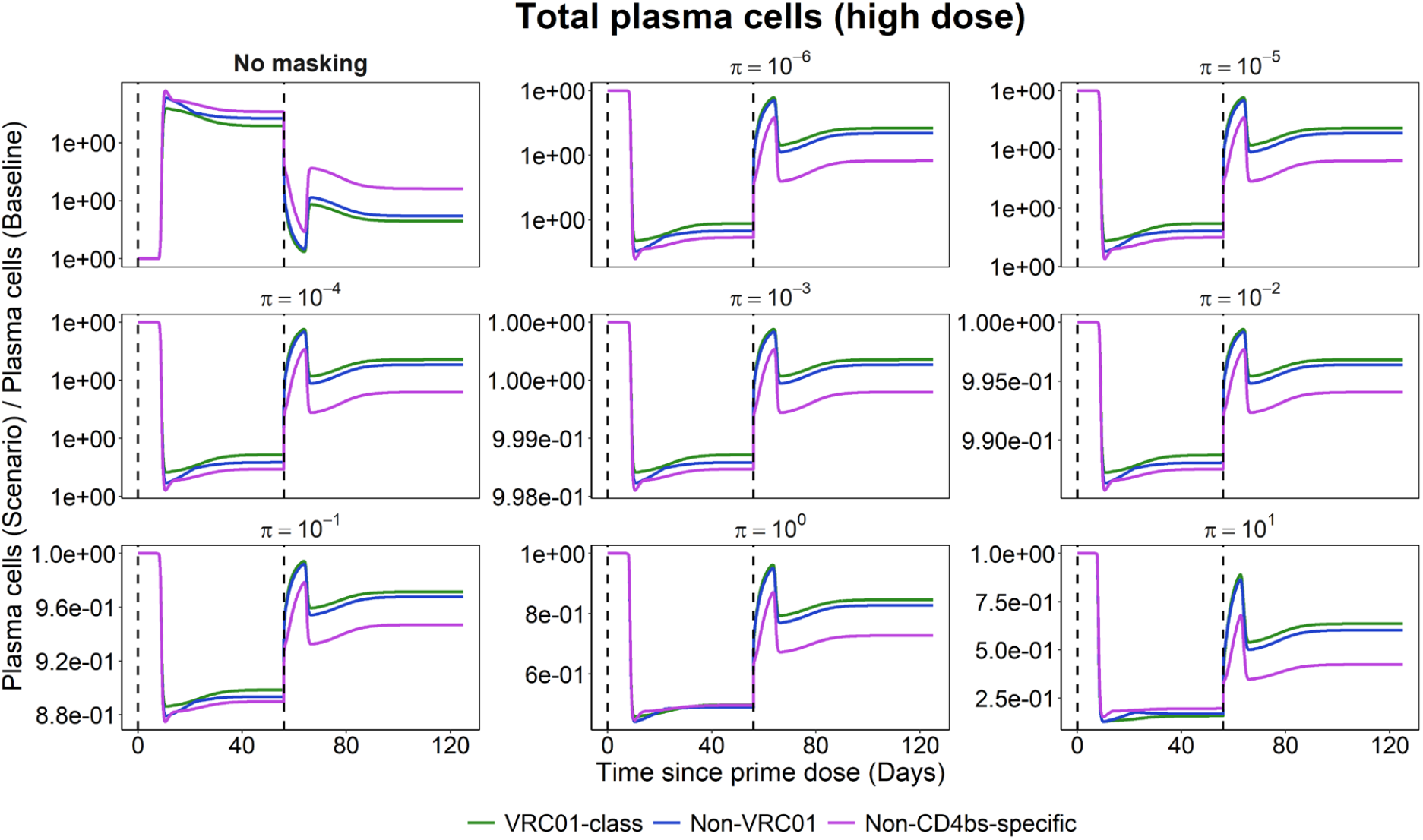
Effect of antibody masking on vaccine-induced plasma-cell dynamics. Ratio of plasma cells for different values of the immunogen–antibody binding rate (*π*) relative to the plasma cells at *π* = 10^−7^ (baseline).

**Figure SM13:**
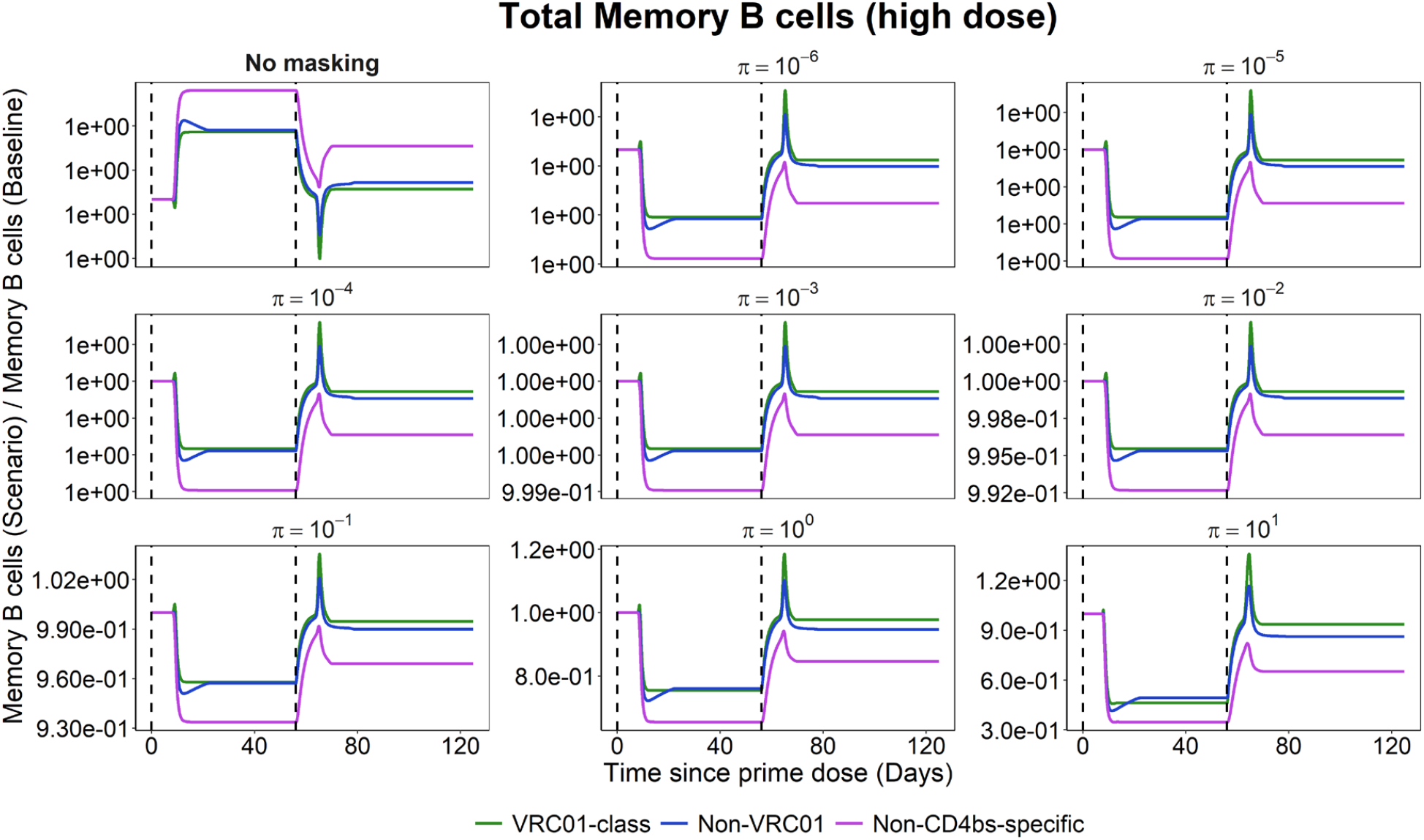
Effect of antibody masking on vaccine-induced memory B-cell dynamics. Ratio of memory B cells for different values of the immunogen–antibody binding rate (*π*) relative to the memory B cells at *π* = 10^−7^ (baseline).

**Figure SM14:**
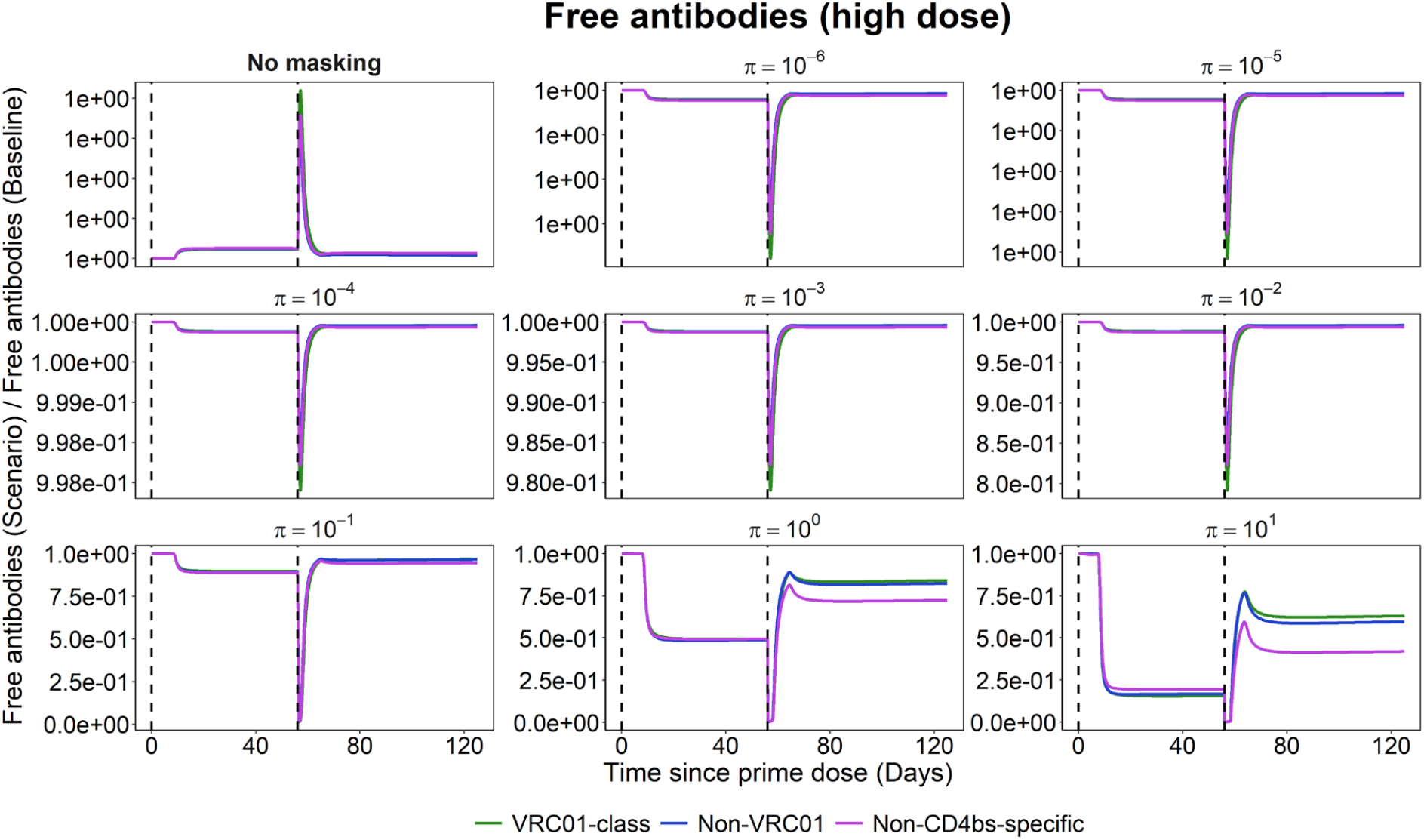
Effect of antibody masking on vaccine-induced antibody dynamics. Ratio of antibody levels for different values of the immunogen–antibody binding rate (*π*) relative to the antibody level at *π* = 10^−7^ (baseline).

**Table SM2.** Estimated population parameters across fixed antibody-immunogen binding rates. Values are population estimates (RSE, %). Each column corresponds to a model fitted with log_10_(*π*) fixed at the indicated value. Parameters prefixed with log_10_ were estimated on the logarithmic scale; RSEs are reported on the same scale. NaN indicates that the RSE could not be evaluated.

| Parameter | $\pi = 10^{-7}$ | $\pi = 10^{-1}$ | $\pi = 1$ | $\pi = 10$ |
| --- | --- | --- | --- | --- |
| $\theta_a$ | $1.2 \times 10^{-3}$ (126) | $7.1 \times 10^{-5}$ (87.1) | $9.8 \times 10^{-5}$ (NaN) | $1.4 \times 10^{-4}$ (39.3) |
| $\theta_b$ | 0.023 (162) | 0.36 (26.7) | 0.29 (23.3) | 0.19 (53.4) |
| $\log_{10}(\zeta_m)$ | 1.96 (1.04) | 2.11 (0.317) | 2.05 (0.532) | 2.09 (0.766) |
| $\log_{10}(\gamma_n)$ | -11.03 (1.01) | -10.99 (1.12) | -10.79 (2.61) | -10.24 (1.98) |
| $\log_{10}(\gamma_{m1})$ | -0.85 (5.08) | -0.91 (0.473) | -0.82 (2.37) | -0.83 (3.36) |
| $\log_{10}(\gamma_{m2})$ | -0.33 (19.5) | -0.63 (NaN) | -0.54 (4.50) | -0.54 (6.61) |
| $\log_{10}(\gamma_{m3})$ | -0.14 (35.1) | -0.17 (11.2) | -0.089 (32.6) | -0.058 (51.7) |
| $\delta_M$ | 0.026 (8.72) | 0.027 (8.58) | 0.026 (8.64) | 0.027 (8.64) |
| $\delta_l$ | $8.0 \times 10^{-4}$ (NaN) | $6.9 \times 10^{-4}$ (34.6) | $8.1 \times 10^{-4}$ (NaN) | $1.0 \times 10^{-3}$ (NaN) |
| $\log_{10}(K_{g1})$ | 7.81 (6.45) | 6.84 (6.64) | 7.09 (10.3) | 6.54 (8.49) |
| $\log_{10}(K_{g2})$ | 3.97 (NaN) | 5.67 (14.5) | 5.52 (47.0) | 3.02 (75.4) |
| $\log_{10}(K_{g3})$ | 7.96 (25.5) | 7.81 (7.15) | 8.19 (16.9) | 8.22 (10.6) |
| $\log_{10}(\delta_{11})$ | -2.39 (6.04) | -2.31 (3.31) | -2.39 (4.33) | -2.44 (3.42) |
| $\log_{10}(\delta_{12})$ | -3.45 (5.78) | -2.96 (8.17) | -3.03 (20.7) | -3.08 (14.2) |
| $\log_{10}(\delta_{22})$ | -1.58 (7.19) | -1.43 (5.33) | -1.47 (6.19) | -1.57 (5.38) |
| $\log_{10}(\delta_{13})$ | -3.19 (32.4) | -3.07 (8.47) | -3.28 (15.6) | -3.23 (11.3) |
| $\log_{10}(\delta_{23})$ | -2.34 (28.1) | -2.28 (13.0) | -2.44 (20.7) | -2.28 (15.2) |
| $\log_{10}(\delta_{33})$ | -1.08 (11.3) | -0.93 (7.53) | -1.01 (9.95) | -1.04 (7.73) |
| $\log_{10}(\Psi)$ | 16.55 (0.574) | 16.56 (0.348) | 16.55 (0.607) | 16.48 (0.362) |

